# Niche-dependent epistasis within the cancer genome shapes metastatic fitness

**DOI:** 10.64898/2026.10.02.756296

**Authors:** Xiyu Liu, Le He, Min Zhao, Samantha M. Golomb, Wei Ma, Xinrui Yang, Emilija Aleksandrovic, Hazel M. Borges, Kevin M. Dean, Lei Wang, Gary C. Hon, Dailin Gan, Hao Zhu, Jun Li, Lin Xu, Siyuan Zhang

## Abstract

Epistasis, the genetic interaction between loci, directs non-additive genotype-to-phenotype outcomes. Within a multicellular organism, how the tissue niche (E) shapes gene-by-gene epistasis (G×G) and its phenotypic consequences remains poorly studied. Using tumor lung metastasis as an experimental model, we mapped G×G×E through combinatorial CRISPR perturbation coupled with depletion of defined host myeloid populations in mice. The metastatic niche first selects disseminated tumor cells on G×G epistatic fitness, then drives transcriptome convergence regardless of genotype. Removing anatomically distinct myeloid populations reshaped the metastatic epistasis landscape despite identical tumor genotypes. Gene-by-gene-by-environment epistasis therefore operates at the organismal level, with interaction magnitude and sign set by niche cells. Beyond metastasis, our data suggest that the tissue niche conditions how genetic interactions translate into divergent phenotypes.

## Introduction

Epistasis provides a foundational framework for understanding how genetic effects manifest through cellular genetic or environmental contexts (*1, 2*). Originally coined in classical genetics to describe how one genetic locus masks or modifies the phenotypic effect of another (*3*), the concept has been expanded substantially across modern biology: in quantitative genetics, epistasis describes departures from additivity of genetic loci contributing to trait variation (*1, 4–8*), while in systems biology, epistasis broadly refers to pathway organization, relationships, and cellular wiring along natural developmental trajectories or in response to experimental perturbations (*9–15*). Importantly, genetic interactions are not necessarily fixed properties of gene pairs but can be extensively rewired by DNA-damaging agent, establishing the concept of differential epistasis in response to genotoxic stress (*16*).

The concept of idiosyncratic epistasis is particularly relevant in cancer, where tumor cells exhibit profound genetic heterogeneity and phenotypic diversity contingent on cancer genome mutational background (*17–20*). The oncogenic effects of extensive, seemingly stochastic tumor genomic alterations are collectively shaped by co-occurring genetic alterations, epigenetic cell state rewiring along tumor progression trajectories, and adaptive responses to anti-cancer therapies (*21–24*). Non-additive dependencies can also expose therapeutically actionable synthetic-lethal vulnerabilities in cancer (*25–27*). Metastasis, where disseminated tumor cells (DTCs) interact with the metastatic niche environment, represents the most consequential yet least tractable manifestations of epistasis (*21, 28–31*). Recent advances in combinatorial perturbation and functional genomics now allow epistasis to be interrogated systematically across diverse biological systems, including mammalian cells (*15, 32–38*). Yet, the principles governing multigenic epistatic gene-gene interactions (GIs) during metastatic progression at the metastatic niche, where tumor-intrinsic and niche-associated factors intersect, remain largely unknown.

Here, we conducted epistatic GI analysis using an experimental lung metastasis model by coupling multiplexed in vivo dual-gene perturbation (G×G) with single-cell transcriptomics and with genetic depletion of metastatic niche (E) cells. At the temporal scale, the success of early metastatic colonization is selective and largely driven by G×G×E epistatic GI at the metastatic niche. At single-cell resolution, however, the transcriptomes of surviving metastatic cells converged in vivo regardless of genotype, indicating that an epistatic interaction acts on which genotypes survive the niche rather than on the transcriptional state those survivors adopt.

Depleting a single host myeloid population changed the sign or magnitude of most measured genetic interactions, and depleting two anatomically distinct lung macrophage populations produced different interaction maps. These observations reframe metastasis as a two-layer selective and adaptive process and establish metastatic epistasis as a property of the tumor genome together with the host niche (G × G × E), driving metastatic colonization and fitness.

## Results

### Niche-associated co-occurring genome dysregulation in clinical metastases

The dysregulated cancer genome is extremely heterogeneous, harboring co-occurring or mutually exclusive CNVs and mutations, and transcriptomal plasticity (*21*). Observed tumor genome heterogeneity is not an exclusive attribute of tumor cell-intrinsic property. Rather, such seemingly chaotic, potentially epistatic, G × G interactions residing in the tumor genome are likely to be modulated and conditioned by the tumor microenvironment (niche) to manifest as tumor fitness outcomes - reminiscent of the gene-by-gene-by-microenvironment (G × G × E) model in classical genetic epistasis studies (*21*). Whether the metastatic microenvironment conditions the epistatic architecture of the tumor genome, altering the magnitude and even the sign of driver interactions, is unresolved in the field (**Fig. 1A**).

**Fig. 1.**
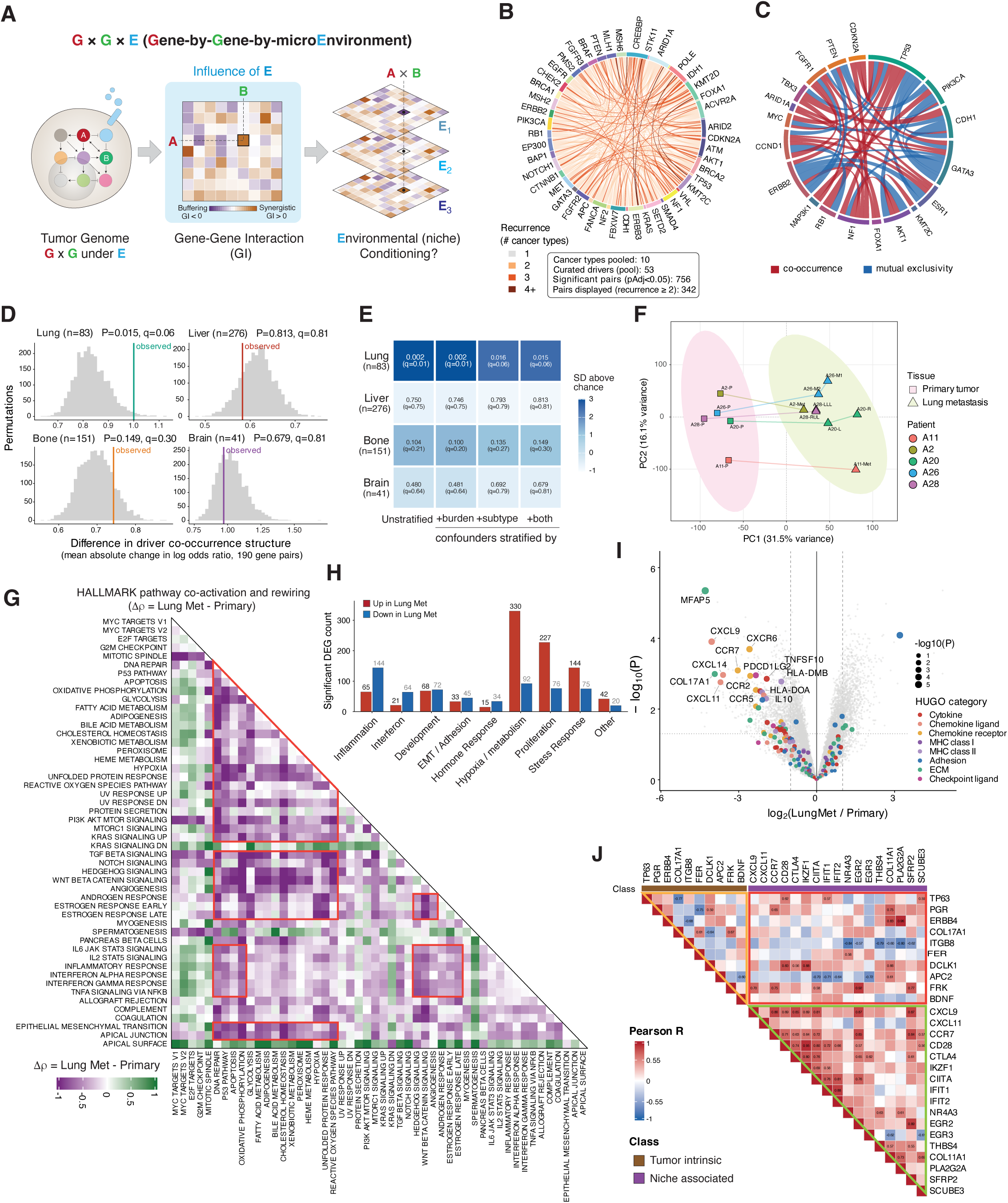
Niche-associated co-occurring genome dysregulation in clinical metastases. (A) Schematic of the central question: how is the fitness consequence of a co-occurring tumor genome dysregulation influenced by the tumor microenvironment (G × G × E). (B) Driver gene co-occurrence is pervasive across metastatic cancers (53 curated drivers, 10 TCGA cohorts); chords, pooled significant pairs (adjusted P < 0.05); colour, number of cancer types in which the pair recurs. (C) Co-occurrence architecture of breast distant metastases (MSK-MET, n = 1,512; 20 drivers); red, co-occurrence; blue, mutual exclusivity (Fisher exact, FDR < 0.10); width, |log odds ratio|. (D) Differential driver co-occurrence structure analysis of metastasis-derived specimens. Grey, matched null; colored line, observed. (E) Co-occurrence structure analysis across four confounder models. Permutation P with the Benjamini–Hochberg q across the four sites; fill, SD above the matched null (random chance). (F) Paired primary and lung-metastasis transcriptomes separate along PC1 (GSE110590; 5 patients, 13 samples). Each point represents an independent tumor specimen. Points with the same color are specimens from the same patient, and connecting lines link the matched primary tumor–lung metastasis pair from each patient. (G) Heatmap of hallmark pathway co-activation rewired across the primary-to-lung-metastasis transition (Δρ = lung metastasis − primary). Each cell represents the difference in Spearman correlation between two pathway activity scores. The upper and lower triangles display the same symmetric pairwise correlation-difference matrix; diagonal cells represent self-correlations. Green indicates increased pathway co-activation in lung metastases, purple indicates decreased co-activation, and white indicates little or no change. Red outlines highlight representative pathway blocks discussed in the text; the direction and magnitude of each change are encoded exclusively by the cell color and color scale. (H) Significant differentially expressed genes per Hallmark module (paired limma; adjusted P < 0.05, |log2 FC| > 1; n = 5), colored by direction in lung metastasis. (I) Volcano plot of differentially expressed niche associated genes in lung metastasis. (J) Gene-level co-activation of tumor-intrinsic and niche-associated factors in lung metastases (Pearson r; n = 8).

We first asked whether pairwise structure was sufficiently recurrent in TCGA pan-cancer dataset. Across 10 TCGA PanCancer Atlas cohorts, 53 consensus cancer drivers formed 756 significant co-occurring pairs, 342 of which recurred in at least two tumor types (**Fig. 1B**). In MSK-MET breast distant-metastatic cohort (n = 1,512 distant-metastatic patients), 20 recurrently altered canonical drivers formed 44 significant associations with 24 co-occurring and 20 mutually exclusive (*FDR < 0.10*) (**Fig. 1C**). We next asked whether this architecture differs by metastatic site (niche). For each co-altered pair, we compute the mean absolute difference in Haldane-corrected pairwise log-odds ratio across all 190 gene pairs and evaluated it against a permutation null matched on intrinsic subtype and metastatic-burden. Lung metastases, but not liver, bone and brain, showed a significant separation from null (*p = 0.015, q = 0.06*). (**Fig. 1D, fig. S1A-D**), regardless of whether tumor subtype and burden were controlled singly, together, or not at all (**Fig. 1E**). In addition to co-occurrence of genomic alterations, we next asked whether lung metastases adopt unique co-occurrent transcriptome patterns compared with paired primary tumors. We analyzed GSE110590, a paired primary breast tumor versus lung-metastasis RNA-sequencing (RNA-seq) series. In the paired subset (5 patients, 13 samples: 5 primary and 8 lung metastases), principal-component analysis separated samples primarily by tissue (PC1, 31.5% of variance) rather than by patient (PC2, 16.1%) (**Fig. 1F**). The co-regulatory relationships among Hallmark pathways were extensively rewired during lung colonization: the co-activation modules, tightly coherent in primary tumors, fragmented into smaller and partially anticorrelated sub-modules in lung metastases (**fig. S1E**), and the differential (Δρ = ρ in lung metastasis minus ρ in primary, across 50 gene sets) was most pronounced among developmental-signaling (TGF-β, Notch, Hedgehog, WNT-β-catenin, angiogenesis), hormone-response, and inflammatory/interferon modules (**Fig. 1G**). Paired differential expression analysis identified predominantly higher hypoxia/metabolism, proliferation, and stress-response gene expression in lung metastases (330 of 422, 227 of 303, and 144 of 219 significant genes, respectively), and predominantly lower inflammation, developmental-signaling, and hormone-response expression (144 of 209, 64 of 85, and 34 of 49) (**Fig. 1H**). Interestingly, extracellular-matrix gene *MFAP5*, chemokines (*CXCL9, CXCL11, CXCL14*), chemokine receptors (*CCR2, CCR5, CCR7, CXCR6*), MHC class II genes (*HLA-DMB, HLA-DOA*), checkpoint ligands (*PDCD1LG2* and *IL10*) were predominantly lower in lung metastases – suggesting a lung niche-immune engagement (**Fig. 1I**). We then asked whether these niche-associated factors were co-active with tumor-intrinsic programs across lung metastases. DawnRank scores, which diffuse each sample’s differential expression over a gene interaction network (*39*) and are used here without filtering for DNA alteration, co-varied within and between the two gene classes across the eight lung metastases (77 of 351 pairs at |r| ≥ 0.55, a descriptive threshold; **Fig. 1J**). The correlated pairs nominate specific intrinsic-by-niche combinations for combinatorial perturbation functional study (**Fig. 2A**). The correlation matrix was not uniform but resolved into three blocks with distinct signatures. Niche × niche pairs were predominantly and coherently positively correlated, forming a clustered block of high positive correlation while intrinsic × intrinsic pairs, by contrast, contained markedly weaker or negative correlations. Interestingly, intrinsic × niche pairs showed more diverse correlation patterns (**Fig. 1J**). This candidate set formed the basis of a combinatorial library designed to test whether tumor-intrinsic and niche-associated perturbations jointly alter metastatic fitness in vivo (**Fig. 2A**).

**Fig. 2.**
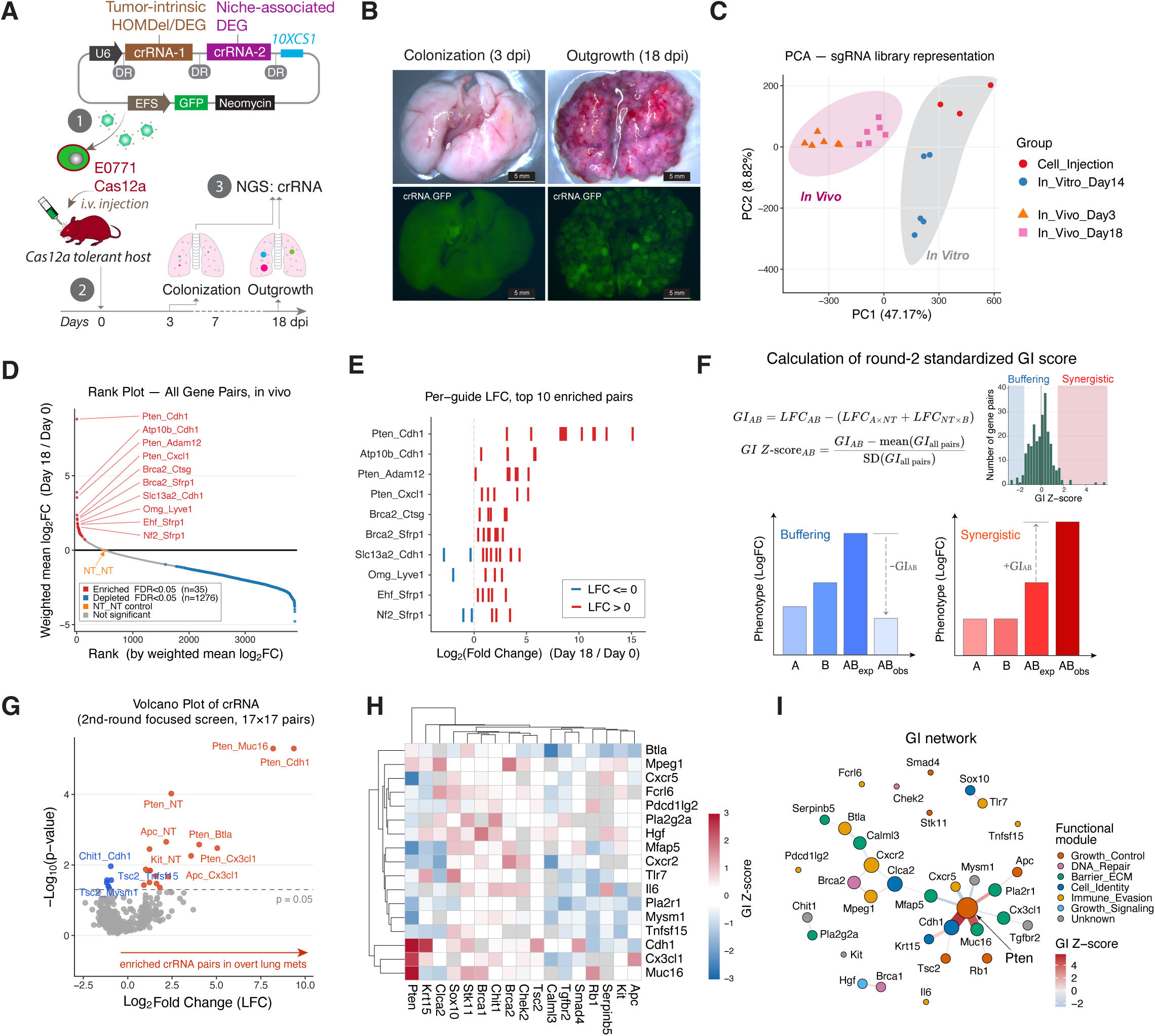
Epistasis GI network governs metastatic gene interactions. (A) In vivo combinatorial dual-Cas12a screen: a lentivirus expresses paired crRNAs against one tumor-intrinsic and one niche-associated gene (91 × 78 grid, 169 genes, 7,098 pairs), delivered into E0771 cells carrying doxycycline-inducible AsCas12a, injected intravenously into Cas12a-tolerant hosts and recovered from the lungs at 3 and 18 dpi. (B) Representative lungs at 3 and 18 dpi; brightfield (top), GFP (bottom). Scale bars, 5 mm. (C) PCA of construct-level counts across 20 samples (injected pool, n = 3 biological replicates; in vitro day 14, n = 5; lungs at 3 and 18 dpi, n = 6 each). Ellipses, 80% normal-theory. (D) Rank plot of 3,886 first-round crRNA pairs (3,716 dual-knockout, 169 single-knockout controls, one pooled NT×NT reference) by NT×NT-corrected weighted-mean log2 fold change (18 dpi versus the injected pool). (E) Per-guide log2 fold change versus the injected pool for the ten most enriched pairs; red, positive; blue, negative. (F) Genetic-interaction score calculation used in (H) and (I). For each pair the interaction effect is the observed pair log2 fold change minus the sum of the single-gene log2 fold changes against non-targeting partners, centered and divided by the standard deviation across the 252 tested pairs. Histogram, the observed scores; shading marks |GI Z| > 1.5, the threshold used to draw an edge in (I). In bar charts schematic, A and B denote the single-knockout measurements against NT partners, AB_exp = sum, and AB_obs = observed double knockout. (G) Volcano plot of the second-round focused 17 × 17 screen (286 constructs: 252 gene × gene pairs and 34 single-knockout controls); dashed line, p = 0.05. (H) GI Z-score heatmap for the second round (252 of 289 designed pairs); symmetric scale ±3, zero at white; rows and columns clustered by average linkage on the centred GI Z-score. (I) Strong-edge GI network at |GI Z| > 1.5, 23 edges among 26 of the 34 genes; nodes coloured by curated functional module.

### Epistasis GI network governs metastatic gene interactions

To test the functional significance of co-occurrent genetic events observed in clinical metastases in determining metastatic fitness at the lung metastatic niche, we conducted an *in vivo* Cas12a CRISPR-based combinatorial dual-knockout (KO) screen using syngeneic E0771 breast-cancer experimental lung metastases model. We broadly defined a tumor-intrinsic gene set and a niche-associated gene set based on co-occurrent genetic events (both DNA and RNA level, **Fig. 1**, and **table S1**) and designed a custom Cas12a CRISPR RNA (crRNA) screen library (round 1, **table S2**) where each crRNA construct represent a unique pair of a tumor-intrinsic gene (crRNA-1; co-currently deleted or downregulated in lung metastases) with a niche-associated gene (crRNA-2; microenvironment, immune or extracellular-matrix etc.), pairing 91 tumor-intrinsic against 78 niche-associated genes (169 genes; 7,098 designed combinations, **table S3**).

To mitigate potential adaptive immune rejection of Cas12a-expressing tumor cells, we generated whole-body Rosa26-Cas12a expressing transgenic mice as Cas12a immunotolerant hosts for in vivo metastases screen (**fig. S2A-F**). Briefly, we used CRISPR/Cas9 mediated Rosa26-knock-in approach (*40*) to generate Cas12a-mCherry line (LSL-Cas12a), followed by crossing with a CMV-Cre line to generate offspring (Rosa26-Cas12a) in which both bone-marrow cell and brain tissue showed the expected spread mCherry expression (**fig. S2A-C**). To validate Cas12a function, we isolated primary astrocytes of Rosa26-Cas12a mice and transduced them with a lentivirus encoding a crRNA array targeting Trp53, Pten, Rb1 (**fig. S2D**). Amplicon sequencing confirmed 70-80% gene editing efficiency (n = 3 independent replicates per locus) with reproducible indel spectra at all three target loci (**fig. S2E-F**).

We conducted the first round of in vivo functional epistatic GI screen using experimental lung metastasis model (**fig. S2G**). Cas12a-expressing E0771 cells were transduced with a lentiviral crRNA screening library (**fig. S2H-I)** and subsequently injected intravenously (i.v.) into Cas12a-tolerant hosts. Amplicon sequencing confirmed broad gene-pair representation of the first-round dual-crRNA library in the plasmid pool, with 122,592 of 233,684 designed constructs (52.5%) detected at ≥3 reads, collectively covering 7,097 of 7,098 designed gene pairs (**fig. S2H**).

Profiling of the transduced E0771 cell pool across three independent biological replicates revealed increased construct-level representation skew while retaining broad gene-pair coverage, providing the baseline for quantifying subsequent selection during metastatic colonization (**fig. S2I**). The dual crRNA cassettes were recovered from in vivo lung tissue at early colonization (3 dpi, **Fig. 2B**) and metastatic outgrowth stage (18 dpi, **Fig. 2B**) and analyzed via amplicon sequencing, alongside a parallel culture of the same pool of library-infected cells maintained in vitro for 18 days (See *Methods* for details). Principal-component analysis of crRNA counts distribution across all 20 sequenced samples showed a clear shift of in vivo crRNA library representation from the injected cell pool and the in vitro cultures (**Fig. 2C**), suggesting metastatic selection of disseminated tumor cells in vivo. 3,886 crRNA pairs were detected in vivo and scored, comprising 3,716 of the 7,098 designed dual-knockout combinations, 169 single-knockout controls, and a pooled non-targeting (NT) control. Ranking these by NT-corrected weighted mean log2 fold (LFC) change against the injected pool resulted in 1276 depleted and 35 enriched pairs at BH-FDR < 0.05, with depleted pairs reaching a median LFC of −2.19 (**Fig. 2D**). For the ten most enriched pairs, independent guides targeting the same pair gave concordant fold changes (**Fig. 2E**). Interestingly, knockout of either *Pten* or *Cdh1* alone was neutral or mildly depleting (*Pten*, -0.81, FDR 0.27; *Cdh1*, −1.77, FDR 4.5×10⁻¹⁰), whereas the double knockout was the most enriched pair in the screen at +8.80 (FDR 3.5×10⁻¹⁵) (**Fig. 2D**), suggesting potential epistatic GI between *Pten* and *Cdh1*.

Given the breadth of the first-round library and uneven construct representation after tumor-cell transduction (**fig. S2H–I**), we used this screen as a discovery-stage survey rather than requiring uniform quantitative coverage across all gene pairs. We therefore designed a focused second-round library comprising 17 tumor-intrinsic and 17 niche-associated genes (**table S3**), prioritizing genes represented by detectable constructs in the first-round screen and further informed by their clinical alteration patterns, biological relevance to metastatic fitness and tumor–niche interactions, and suggestive interaction behavior in the discovery screen. This focused design enabled substantially deeper per-construct coverage across 289 gene pairs under matched in vivo conditions (**fig. S2J–K, table S4**). Compared with cell baseline (Day 0), in vivo selection showed an evident redistribution cell populations: median LFC moved from +0.35 at day 3 to −0.51 at day 18 (**fig. S2L-M**). To calculate epistatic GI, for each gene pair, epistasis was defined as the difference between the observed double-knockout LFC and the sum of the two single-gene LFC changes measured with NT partners, with positive GI z-score values indicating synergy (positive) or buffering (negative) phenotypic outcomes (**Fig. 2F**). 17 pairs were enriched in overt metastases at Day 18 compared with cell baseline while 6 pairs depleted (nominal p < 0.05) with Pten-containing pairs exhibited high levels of enrichment, ranging +9.36 with *Cdh1* and +8.19 with *Muc16*, followed by +5.05 with *Cx3cl1* and +4.03 with *Btla* (**Fig. 2G**). In contrast, the same library cultured in vitro for 14 days yielded a single enriched pair, *Pten-Cdh1* itself, at +2.14 (BH-FDR = 0.039), compared with +8.80 in vivo, a difference of 6.67 log2 units (**fig. S2N**). Thus, although the interaction retained a modest cell-intrinsic component, its substantially greater magnitude in vivo indicates strong potentiation by the metastatic niche. The pairwise epistatic interaction matrix heatmap, rendered on a GI sign symmetric scale, showed a mixed GI sign with 138 positive and 114 negative GI interaction pairs with a distinct hierarchical clustering grouped between *Pten* and its co-deletion partner *Cdh1* (GI=+7.63), *Cx3cl1* (GI=+3.43), and *Muc16* (GI=+6.50) (**Fig. 2H and fig. S2O**). To map the topology of the strongest interactions, we constructed a GI network in which edges connect gene pairs when |GI Z| > 1.5 (**Fig. 2I**). Twenty-three of the 252 assayed pairs exceeded this threshold (13 synergistic, 10 buffering), connecting 26 of the 34 genes. *Pten* carried seven strong partners, more than twice the degree of the next-ranked genes (*Cdh1, Clca2* and *Cxcr2*, three partners each). *Pten-Cdh1* synergy was the strongest edge in the network (GI Z = +5.70), consistent with its leading enrichment in the primary screen. In comparison, classic STRING protein-interaction database identified 30 cross-program edges among the same 34 genes (**fig. S2P**), yet only one edge (Pten-Cdh1) overlapped with the genetic interaction network, indicating that nearly all strong epistatic interactions are orthogonal to known physical or functional protein association. We next aggregated GI pairs by curated functional module (**fig. S2Q**). Interestingly, synergistic GIs showed functional annotation enriched with cell identity, growth, DNA repair modules, whereas the most strongly buffering combinations all involve barrier and/or extracellular matrix (ECM) functions (**fig. S2Q**).

### Metastatic Niche Selection Determines Pten-partner Epistasis Outcome

Topology of the GI network highlighted Pten as a central hub, while the identity of the GI partner determined whether its interaction was synergistic or buffering (**Fig. 2I**). To test whether GI sign predicts metastatic outcome, we generated isogenic lines carrying deletion of each of four Pten partners in a Pten-KO background (**fig. S3A-C**) and assessed their impact on the metastatic outcome in independent hosts (**Fig. 3A**). Two partners were drawn from the top-ranked synergistic GI pairs in the screen (Cdh1, Cx3cl1) and one from the buffering extreme (Cxcr5). We also included Tlr7, a weakly buffering partner, because tumoral Tlr7 has been reported to promote breast cancer metastasis through PI3K-AKT, the pathway Pten restrains (*41*). GI sign predicted phenotypic direction in all cases: compared with Pten KO alone, dual knockout of Pten with Cdh1 or Cx3cl1 raised lung burden from 10.1% to 18.9% and 14.8% (*p = 0.041, p = 0.038*), while dual knockout with Cxcr5 or Tlr7 lowered metastatic burden to 4.3% and 3.2% (*p = 0.013, p = 0.0024*) - a sixfold spread across genotypes despite sharing the same tumor-suppressor Pten deletion **(table S**5).

**Fig. 3.**
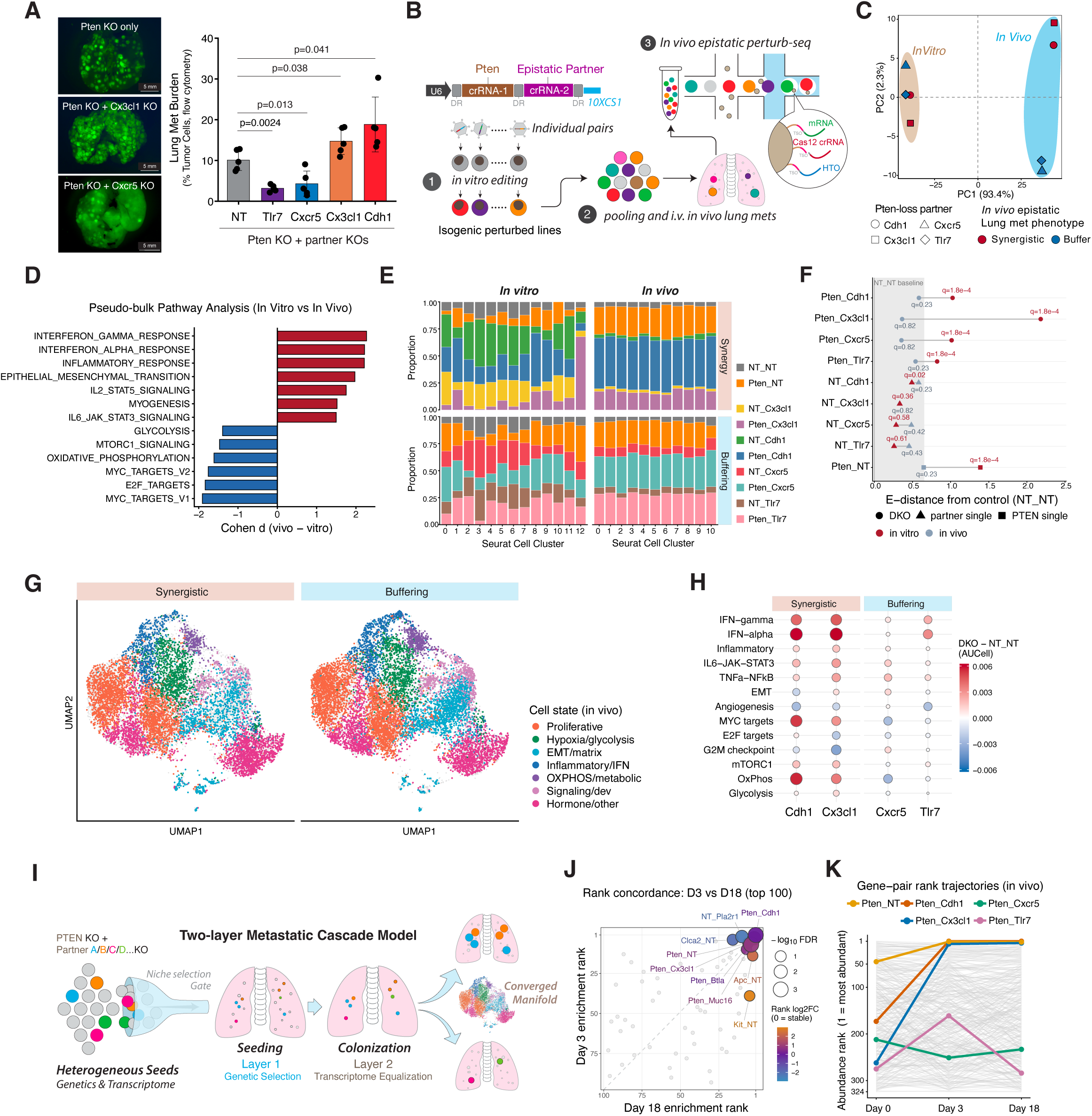
Metastatic niche selection determines Pten-partner epistasis outcome. (A) Lung metastatic burden after intravenous injection of E0771 cells carrying Pten KO with a non-targeting control (NT) or the indicated partner knockout; whole-lung GFP at left. Each point is one mouse (n = 5); bars, mean ± s.d.; p, unpaired two-tailed t-test versus Pten + NT. (B) Screen design. A dual-crRNA enAsCas12a vector targets Pten and the partner from one U6 array. Edited pairs are pooled and injected intravenously; Perturb-seq of tumor cells from lung metastases reads transcriptome, crRNA and mouse of origin (HTO) per cell. (C) Pseudobulk principal-component analysis of the eight double-knockout genotypes. Shape, partner; fill, in vivo phenotype class. (D) Hallmark programs separating the NT_NT control in vivo from the same control in vitro, ranked by Cohen’s d. (E) Cluster composition as P(guide | cluster), by pool and environment. (F) Energy distance from the same-lane NT_NT control in 30 principal components at matched cell number; labels, exact-permutation BH q. Band, control-versus-control 95th percentile. (G) In vivo UMAP split by pool, colored by the seven annotated cell states; gray, all cells. (H) Per-mouse residual, each double knockout minus the NT_NT control of its own mouse (AUCell), over 13 pre-specified niche pathways. Fill, mean over mice; size, absolute mean. (I) Two-layer model: the niche selects among genetic combinations at seeding (Layer 1); survivors converge onto a shared state through colonization and outgrowth (Layer 2). (J) Day 3 versus Day 18 enrichment rank for pairs in the top 100 at both time points. Size, −log₁₀ FDR; fill, rank log₂ fold-change (purple, stable; blue, Day 3; orange, Day 18). (K) Pool-abundance rank of all 324 pairs at Day 0, 3 and 18 (gray), five highlighted.

To resolve the cellular states and transcriptional programs that potentially drive these opposing signs of epistasis during metastasis, we next performed in vivo and in vitro single cell Perturb-seq on the focused Pten-anchored GI panel. We i.v. injected a pool of the isogenic tumor cell lines carrying the phenotype-validated Pten-partner perturbations described above to establish in vivo lung metastases. Both the cell-line pool and the metastatic tumor cells recovered from the lungs were subjected to multimodal single-cell sequencing, with simultaneous capture of mRNA, Cas12a crRNAs and hashtag oligonucleotides (HTOs), enabling perturbation-resolved characterization of transcriptional states associated with genetic interactions during metastatic outgrowth.

At the global transcriptome level, PCA analysis showed that PC1, accounting for 93.4% of the total variance, separated cells derived from in vitro from those recovered from lung metastases, whereas perturbation genotype-dependent variation was confined largely to PC2, which explained only 2.3% of the variance (**Fig. 3C**). This pronounced *in vivo* metastatic niche context-dependent reprogramming was further reflected in marked shifts in transcriptional-state composition and cluster-associated pathway programs (**fig. S3D&S3E**). At the Pathway-level, compared with their *in vitro* counterparts, tumor cells recovered from lung metastases showed increased inflammatory and interferon-related programs, whereas proliferative programs, including MYC and E2F signaling, were reduced (**Fig. 3D**).

This dominant metastatic-niche reprogramming markedly overshadowed the effects of individual perturbations on the cellular transcriptome. *In vitro*, each single- and dual-perturbation genotype shifted cell transcriptomes in diverse directions, as evidenced by their uneven distribution across Seurat clusters (**Fig. 3E**) and by the transcriptomic distance (E-distance) induced by each perturbation relative to the NT_NT control (**Fig. 3F**). *In vivo*, however, perturbation effects were largely equalized, as recovered metastatic tumor cells shared similar distributions across Seurat clusters (**Fig. 3E**) and E-distances indistinguishable from control (**Fig. 3F**), suggesting a convergence of cell state regardless of the heterogeneity of perturbation. Differential expression analysis identified a broad niche-adaptation program upon lung colonization (**fig. S3F**). While epistatic GI modelling of the transcriptome manifold suggested a trend in which the primary contribution shifted from *Pten* KO *in vitro* to partner-gene KO *in vivo* (**fig. S3G**), perturbation-induced transcriptomic distances and double-KO-associated global transcriptional signatures were largely diminished *in vivo* (**fig. S3H, I**). Despite this convergence of global cell state imposed by the metastatic niche, some genotype-specific transcriptional effects remained detectable at a finer functional resolution. The *in vivo* transcriptional landscape comprised functionally distinct cell states spanning a broad spectrum of proliferative, inflammatory, metabolic and mesenchymal programs (**Fig. 3G**). AUCell analysis further indicated a residual, directionally consistent genotype effect, with synergistic perturbations deviating further from the NT_NT baseline than buffering perturbations across inflammatory and metabolic programs (**Fig. 3H, fig. S3J**).

The observation that *in vivo* transcriptome equalization overrides the effects of tumor-cell perturbation is counterintuitive, suggesting that the transcriptome alone is not sufficient to explain the sixfold difference in metastatic burden (**Fig. 3A**). One possibility is that the metastatic burden phenotype is decided during seeding (Day 0 to Day 3), when the niche selects which genotypes survive. The shared transcriptional state of the survivors, despite their diverse genotypes of origin, would then reflect a dominant common pattern of adaptation to the metastatic niche during outgrowth, rather than genotype-specific adaptation (**Fig. 3I**). This model predicts that the rank order of genotypes in the lung is set early and preserved thereafter. To test this hypothesis, we reanalyzed the second-round screen (**Fig. 2**) resolved by metastasis stage.

Among pairs enriched at both time points, Day 3 and Day 18 ranks were closely concordant (**Fig. 3J**). Tracking individual pairs from Day 0 through Day 3 to Day 18 revealed a “sorting” event within the seeding window: synergistic pair *Pten_Cdh1* rose from rank 173 to rank 1 within three days and held that position to Day 18, and *Pten_Cx3cl1* rose from 262 to 7 by Day 3 and to 5 by Day 18, whereas the buffering pairs (*Pten_Cxcr5* and *Pten_Tlr7*) remained at low ranks throughout (**Fig. 3K**). Niche selection among genotypes, particularly of the synergistic GI pairs, therefore occurs within the critical seeding-colonization window, rather than arising as a consequence of epistatic GI within the adaptive transcriptome manifold. Once selected, the surviving clones outgrow into overt metastases that share a converged, niche-driven cell state, regardless of differences in their genetic composition.

### Host CX3CR1⁺ Cell Niche Gates Metastatic Epistasis In Vivo

Our results support a two-layer metastatic cascade model (**Fig. 3I**) that elaborates the multistep selective and adaptive nature of the classic seed-and-soil framework, in which the metastatic niche imposes a selection pressure that gates which arriving metastatic tumor cell to survive. Yet the seed-and-soil relationship is dynamic and reciprocal: the niche itself co-evolves, remodeling in response to physiological, pathological, and therapeutic cues, and consequently reshaping which GI it permits. The epistatic GI we observed is therefore unlikely to arise solely from tumor cell-autonomous mechanisms. Instead, it may be realized through engaging co-evolved host cells at the metastatic niche, or through a combination of the two. The metastasis-promoting synergistic GI Pten_Cx3cl1 has suggested this possibility, as the tumor cell presents CX3CL1, whose canonical receptor CX3CR1 is abundantly expressed on myeloid cells positioned at the seed-soil interface (**Fig. 4A**). Recent work further shows that lung-resident alveolar macrophages can regulate the timing of breast cancer metastatic outgrowth (*42*).

**Fig. 4.**
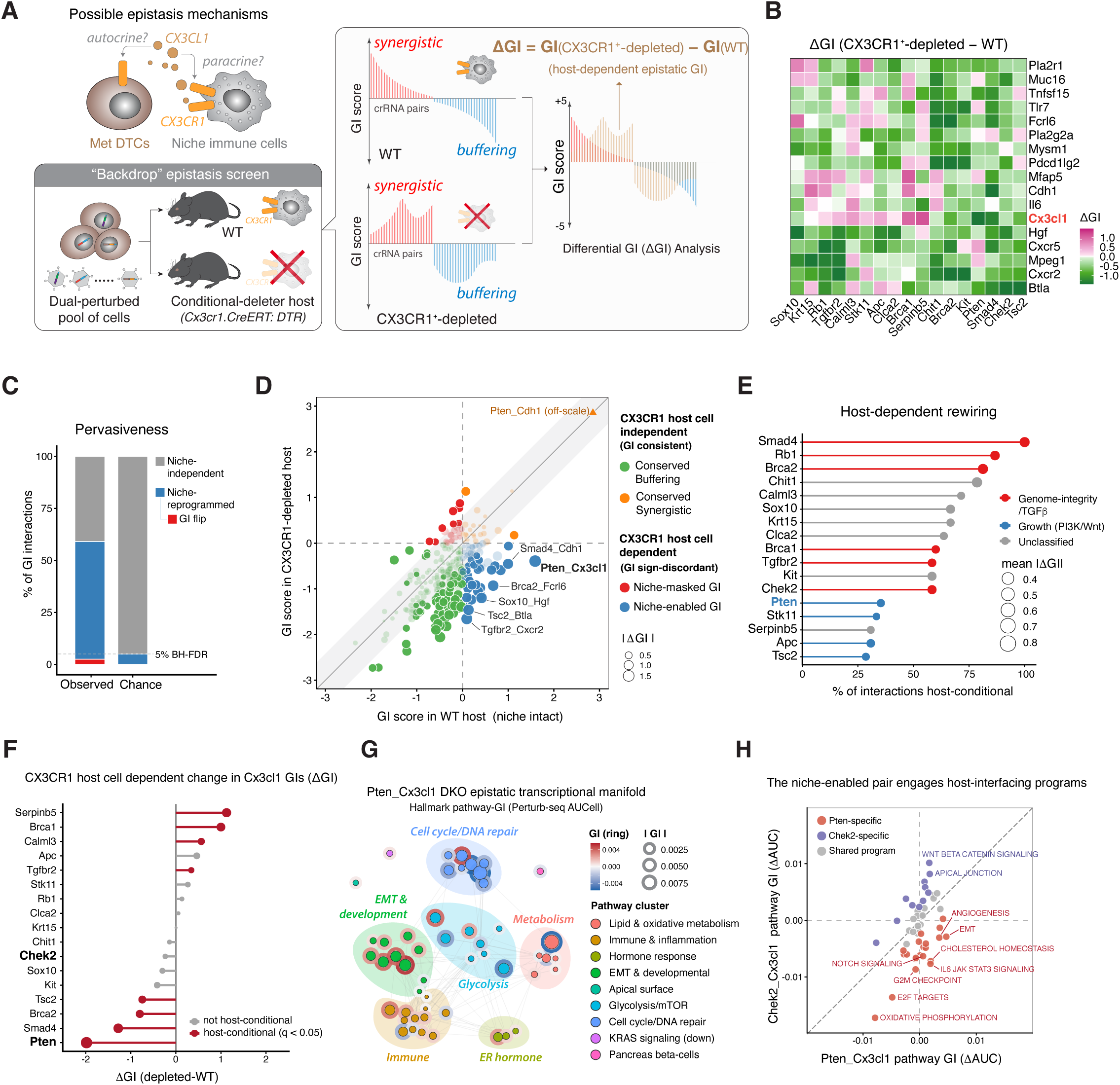
Host CX3CR1⁺ niche gates metastatic epistasis in vivo. (A) Schematic of the host-backdrop epistasis screen. A pooled dual-guide Cas12a library covering all pairwise combinations of 17 tumor drivers and 17 niche/immune partners (289 pairs), plus single-gene and double-non-targeting (NT_NT) controls, was assayed in E0771 cells recovered after metastatic outgrowth in niche-intact (WT) and CX3CR1⁺-depleted hosts (Cx3cr1.CreERT:DTR plus diphtheria toxin; n = 7 mice per arm). GI is the deviation from the additive expectation of the two single perturbations, in log₂ fold-change units; ΔGI = GI(depleted) − GI(WT). GI > 0, synergy; GI < 0, buffering. (B) Heatmap of ΔGI across 17 drivers (x) by 17 partners (y). Diverging scale centered at zero, clipped at the 97th percentile of |ΔGI|; ** q < 0.05, * q < 0.10. (C) Pervasiveness of host-dependent rewiring. (D) Genetic-interaction state space. Each pair is plotted at its GI in the WT (x) and depleted (y) host. Diagonal, host-invariant interactions; shaded band, 95% host-label permutation null (|ΔGI| ≈ 0.58). Fill, sign quadrant (conserved buffering, conserved synergistic, niche-masked, niche-enabled); size, |ΔGI|; opacity, bootstrap significance. n = 289 pairs, of which 198 are genuine interactions (q < 0.05 in ≥ 1 host). (E) Host-conditional rate per driver. (F) CX3CR1-dependent change of ΔGI of each driver’s interaction with Cx3cl1. (G) Hallmark-program network for Pten_Cx3cl1 on a fixed Jaccard similarity scaffold. Ring color, signed pathway-level GI; size, |GI|. (H) Programs by pathway-level GI in Pten_Cx3cl1 (x) versus Chek2_Cx3cl1 (y).

To determine whether changing the host microenvironment is sufficient to rewire the pairwise epistatic GI landscape of successful metastases, we repeated the in vivo dual-crRNA GI screen with the same perturbation library in matched niche-intact (WT) and CX3CR1⁺-myeloid-cell-depleted (Cx3cr1.CreER:DTR) hosts. This niche cell depletion (a “backdrop”) epistasis screen quantifies CX3CR1⁺-myeloid-cell-dependent epistasis directly, as the differential GI (ΔGI) between the two hosts (**Fig. 4A**). ΔGI was calculated as GI in depleted hosts minus GI in WT hosts. For interactions that retain their sign, weakening denotes a decrease in absolute GI and strengthening denotes an increase; interactions that cross zero undergo a sign reversal. We validated the efficient depletion of CX3CR1⁺ monocytes after diphtheria toxin treatment by flow cytometry (**fig. S4A, B**, **table S**6). ΔGI analysis revealed widespread rewiring of the epistatic GI landscape following depletion of CX3CR1⁺ niche cells (**Fig. 4B** and **fig. S4C, D**). This rewiring was pervasive: 117 of 198 genuine interactions (59%) were significantly reprogrammed by niche depletion, far exceeding the 5% false-discovery ceiling implied by the significance threshold (**Fig. 4C**). Next, we compared changes in GI sign (directionality) between the two host backgrounds. Most GIs retained their buffering or synergistic behavior, representing conserved genetic interactions (**Fig. 4D**). A subset instead differed in sign between hosts: some were synergistic only in the intact niche (niche-enabled), other GIs emerged only after removal of CX3CR1⁺ niche cells (niche-masked). Five of these reversals were significant in both hosts and represent the most extreme form of host-dependent rewiring (**fig. S4E-F**).

Next, we asked whether host-conditional rewiring was restricted to particular genes, either the tumor-intrinsic driver at position 1 or the niche-associated partner at position 2. Across the seventeen drivers, host-conditionality ranged from 29% to 100% of a driver’s interactions. Except for *Smad4*, for which all eleven *Smad4*-associated GIs were rewired, most driver gene GIs showed varying degrees of host-conditionality (**Fig. 4E**). *Pten* is among the least host-conditional drivers in the screen: only 6 of its 17 interactions were altered in DTR hosts. Yet, *Pten_Cx3cl1* carried the largest absolute ΔGI of any *Pten* interaction upon CX3CR1⁺ cell depletion, indicating that the *Pten_Cx3cl1* synergistic GI is primarily driven through tumor-immune crosstalk, rather than autocrine or cell-autonomous mechanisms. Further analysis revealed bidirectional changes in *Cx3cl1*-containing GIs after CX3CR1⁺-cell depletion (**Fig. 4F**). The *Pten_Cx3cl1* interaction reversed from synergistic to buffering (GI +1.59 to −0.39; ΔGI = −1.98). Depletion strengthened buffering GIs for *Smad4_Cx3cl1*, *Brca2_Cx3cl1* and *Tsc2_Cx3cl1* (ΔGI −1.28 to −0.75), while weakening buffering GIs for *Serpinb5_Cx3cl1*, *Brca1_Cx3cl1*, *Calml3_Cx3cl1* and *Tgfbr2_Cx3cl1* (ΔGI +1.13 to +0.34).

To explore the molecular basis of the above difference, we compared the Perturb-seq-derived transcriptome manifolds of two Cx3cl1 interactions - *Pten_Cx3cl1* (host-conditional) and *Chek2_Cx3cl1* (host-independent). Pathway-level comparison revealed multifaceted epistatic remodeling across multiple biological programs upon *Pten_Cx3cl1* double knockout (**Fig. 4G**). Compared with the host-independent *Chek2_Cx3cl1*, *Pten_Cx3cl1* preferentially engaged IL6-JAK-STAT3 signaling, cholesterol homeostasis, and EMT pathways, whereas *Chek2_Cx3cl1* exhibited stronger coupling to WNT-β-catenin signaling and apical junction pathways (**Fig. 4H**). Notably, the *Chek2_Cx3cl1* double knockout produced non-additive transcriptional responses in both the early and late estrogen-response programs (**fig. S4G**). Functionally, compared with either single knockout alone, the *Chek2_Cx3cl1* DKO showed enhanced responsiveness to estrogen-pathway modulation (**fig. S4H**). Together, these findings establish host-derived niche cues as active determinants of genetic-interaction architecture, selectively rewiring epistatic programs according to tumor genetic context.

### MRC1⁺ macrophage loss unmasks Pten-Cdh1 metastatic epistasis

Epistasis is not a tumor-cell-autonomous and fixed genetic property. The phenotypic output of GI is a collective result of the interplay between tumor-cell-intrinsic genome context and niche environment. It has been well documented in model organisms that the sign and magnitude of a given GI depend on the conditions under which it is measured (*5, 43, 44*), and that GI networks reorganize substantially when cells are challenged (*16*). We therefore asked whether the GI landscape of DTCs changes when the metastatic niche differs. Our data in Fig. 4 have demonstrated that depletion of CX3CR1⁺ monocyte-derived macrophages reshaped the GI landscape. In addition to monocyte-derived macrophages, the lung harbors MRC1⁺LYVE1⁺ perivascular interstitial macrophages (IMs), a highly specialized subset of lung tissue-resident macrophages (TRMs) that occupies an anatomically distinct compartment from the peribronchovascular CX3CR1⁺ population and guards lung tissue homeostasis (*45*) (**Fig. 5A**). To distinguish the specific effect of IMs on lung metastatic epistasis, we repeated the in vivo crRNA screen with the identical library in Mrc1.CreERT:DTR hosts. Specific depletion of MRC1⁺LYVE1⁺ perivascular IMs while preserving alveolar macrophages (AMs) was confirmed by flow cytometry (**Fig. 5A**, **fig. S5A-B,** and **table S**6).

**Fig. 5.**
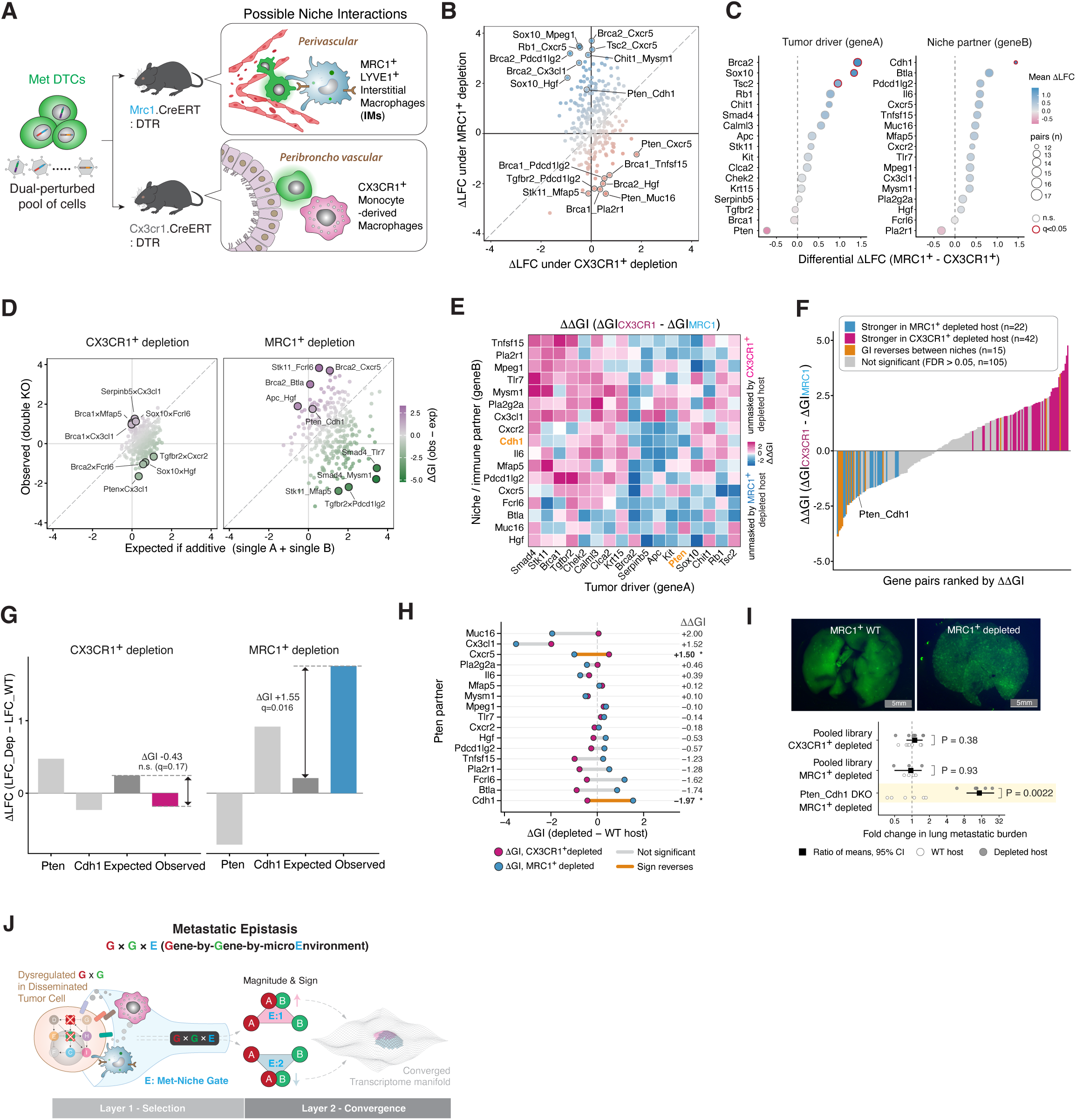
MRC1⁺ macrophage loss unmasks Pten_Cdh1 metastatic epistasis. (A) Design. A dual-perturbed pool of metastatic disseminated tumor cells was screened in parallel in two host-depletion models: Mrc1.CreERT:DTR, depleting MRC1⁺LYVE1⁺ perivascular interstitial macrophages (IMs), and Cx3cr1.CreERT:DTR, depleting CX3CR1⁺ peribronchovascular monocyte-derived macrophages. (B) Per-pair first-order host dependence. ΔLFC (LFC_depleted − LFC_WT) under MRC1⁺ versus CX3CR1⁺ depletion for 269 WT-filtered pairs. (C) Per-gene mean differential ΔLFC (MRC1⁺− CX3CR1⁺), plotted separately for tumor drivers (geneA) and niche partners (geneB). Point size gives the number of contributing pairs; red rings mark q < 0.05. (D) Observed versus expected double-knockout effects for 269 gene pairs passing the wild-type consistency filter, shown separately in CX3CR1-depleted and MRC1-depleted hosts. Each point represents one gene pair. The diagonal line indicates equality between the observed and expected effects. (E) ΔΔGI for all driver × partner combinations, where ΔΔGI = ΔGI_CX3CR1 − ΔGI_MRC1. Pink and blue denote the direction of the between-host difference as indicated on the color bar; the Pten_Cdh1 pair is outlined. (F) All 184 genuine interactions ranked by ΔΔGI and colored by between-host category, with category counts given in the key. Significance is BH-FDR q < 0.05. (G) Single-gene effects, additive expectation and observed double-knockout effect for Pten_Cdh1 in each host. ΔGI and its q value are printed on each panel; the between-host ΔΔGI is given above. (H) All 17 Pten pairs. Each segment joins the interaction’s ΔGI in the CX3CR1⁺-depleted host to its ΔGI in the MRC1⁺-depleted host, colored by between-host category. ΔΔGI is printed at right; * marks q < 0.05. (I) Representative GFP⁺ lung metastases in MRC1⁺ WT and depleted hosts (top; scale bar 5 mm), and fold change in lung metastatic burden, depleted over WT host (bottom). Squares give the ratio of group means with 95% bootstrap confidence interval; circles are individual mice, each divided by the mean of its own assay’s WT group. P values are from exact permutation tests; n = 7 versus 7, 5 versus 3 and 6 versus 6, top to bottom. (J) Model of metastatic epistasis as a Gene-by-Gene-by-microEnvironment (G × G × E) property.

PCA of sample-level crRNA-pair abundance profiles revealed clear separation of the samples by both host-depletion condition (PC2, 11.1% of variance) and screening model (PC1, 54.1%), confirming that both depletions produced a global shift in library composition (**fig. S5C**). Because the two depletion models were generated in independent screening experiments, modest differences were observed between the corresponding LFCs observed in WT hosts. We therefore applied a WT consistency filter, removing the 20 of 289 pairs whose LFCs were opposite in sign across screens with |ΔWT| > 2, thereby excluding pairs whose baseline behavior was inconsistent between screens (**fig. S5D**). Next, we quantified the change in crRNA-pair abundance under different host-cell depletions relative to the matched WT background (ΔLFC, a first-order tumor-burden phenotype readout) and compared these effects between the Mrc1- and Cx3cr1-depletion screens. The resulting pairwise effects diverged substantially between the two host perturbations, with individual genotypes showing preferential responses to either MRC1⁺ or CX3CR1⁺ niche depletion (**Fig. 5B**). We next aggregated these differential effects by gene identity. Only four of 34 anchors showed significant MRC1-biased first-order effects. Three were tumor drivers, *Brca2*, Sox10 and *Tsc2* (q = 0.036 for each). Niche-associated partners showed more restricted effects, with *Cdh1* the only significant partner and its effect stronger in the MRC1⁺ cell-depleted host (mean differential ΔLFC = 1.44, q = 0.00056; **Fig. 5C**). These patterns were robust to per-screen standardization (ρ = 0.90) and were independently recovered by anchor-level ranking (**fig. S5E, S5F**).

We next examined whether these niche-specific dependencies extended to GI relationships (**Fig. 5D**). GI magnitude and distribution differed markedly between the two host-depletion backgrounds. CX3CR1⁺-cell depletion produced a comparatively restricted spectrum of deviations from additivity (SD = 0.79; 8 of 289 pairs with |GI | > 2), whereas MRC1⁺ IM depletion produced substantially stronger positive and negative GIs (SD = 1.61; 103 of 289 pairs). Subtracting each screen’s own WT baseline preserved this contrast, indicating that it does not arise from the different dynamic range of the two screens (ΔGI SD = 0.57 versus 1.49; 0 versus 61 pairs with |ΔGI| > 2; **fig. S5G**). The two niches also remodeled interactions in qualitatively different ways: CX3CR1⁺-cell depletion produced a broader spectrum of GI sign transitions relative to the WT host, whereas MRC1⁺ IM depletion largely preserved GI sign while amplifying its magnitude (**fig. S5H**).

To resolve which niche preferentially conditioned each GI, we calculated ΔΔGI as the difference between the CX3CR1- and MRC1-dependent ΔGI values (**Fig. 5E**). Among 184 gene pairs exhibiting a GI in at least one host context, 79 (43%) showed significantly different ΔGI between the two niches, including 15 whose GI sign reversed between niche contexts (**Fig. 5F**). Aggregating ΔΔGI across each tumor driver’s 17 partners showed that several drivers carry a systematic niche bias, but the spread across partners was broad and most drivers spanned zero, indicating that host dependence is a property of the gene pair rather than of the driver alone (**fig. S5I**). For the *Pten_Cdh1* pair examined below, bootstrap confidence intervals placed ΔGI under MRC1⁺ depletion away from the null (q = 0.016) while ΔGI under CX3CR1⁺ depletion overlapped it (q = 0.168), and the between-host ΔΔGI excluded zero (q = 0.019) (**fig. S5J**), suggesting unique host MRC1⁺-cell dependency.

Among the interactions distinguished by ΔΔGI, we focused on *Pten_Cdh1*. *Cdh1* had emerged as the most niche-biased partner in the first-order layer: it showed the largest difference in ΔLFC between the two hosts and was the only partner-side anchor to reach significance (mean difference = +1.44, q < 0.001) (**Fig. 5C**). In the CX3CR1⁺-depleted host, the *Pten_Cdh1* epistasis residual was indistinguishable from its wild-type value (ΔGI = −0.43, q = 0.17), whereas in the MRC1⁺-depleted host the residual shifted substantially (ΔGI = +1.55, q = 0.016), indicating that loss of MRC1⁺ IMs unmasks the *Pten_Cdh1* interaction (**Fig. 5G**). This host dependence is a property of the interaction partner rather than of *Pten* alone. Among the 17 *Pten* partners examined, only *Cdh1* and *Cxcr5* showed a significant difference in GI between the two depleted hosts, and in opposite directions (ΔΔGI = −1.97 and +1.50, respectively; q < 0.05) (**Fig. 5H**).

To validate the MRC1⁺ IM dependence of the *Pten_Cdh1* interaction, we injected *Pten_Cdh1* DKO cells intravenously into wild-type and MRC1⁺-depleted hosts. Lung metastatic burden increased 14.6-fold in the MRC1⁺ IM-depleted host (*95% CI 8.8–26.2; p = 0.0022*) (**Fig. 5I, table S5**), identifying MRC1⁺ IMs as suppressors of *Pten_Cdh1*-driven metastasis. This effect was specific to the *Pten_Cdh1* genotype rather than a general consequence of removing tumor-suppressive MRC1⁺ cells. Injection of the pooled library into either CX3CR1⁺-depleted or MRC1⁺-depleted hosts left overall lung metastatic burden unchanged, with confidence intervals that exclude any increase beyond 1.5-fold (1.11×, 95% CI 0.80–1.52; 0.95×, 95% CI 0.50–1.46) (**Fig. 5I, table S5**). We next examined the spatial association between MRC1⁺ macrophages and size-matched *Pten*-KO or *Pten_Cdh1* DKO lung lesions in wild-type hosts early in lung colonization (day 3 after injection). Representative sections showed MRC1⁺ IMs closely infiltrated to DKO micrometastases and more sparsely distributed around *Pten*-KO lesions (**fig. S5K**).

Together, these results establish that metastatic epistasis is not a fixed property of the tumor genotype. Epistatic sign, magnitude, and phenotypic outcome are selectively rewired by the surrounding metastatic niche (**Fig. 5J**; G × G × E, gene × gene × microenvironment).

## Discussion

Epistasis is canonically considered a property of interacting gene pairs, yet our findings reveal a fundamental principle of metastatic epistasis at the tumor-niche interface: the metastatic niche can determine both the magnitude and direction of tumor-cell genetic interactions through a two-layer selection in vivo (**Fig. 5J**). Human cancer genomic co-occurrence data provide clinically relevant context for tumor-associated gene epistasis (*21, 46*). However, observational tumor genomic datasets incompletely resolve the cellular and spatial states of the metastatic niche (*47, 48*) and therefore cannot functionally establish G × G × E relationships. Through functional mapping, our study demonstrates niche-directed G × G × E interactions that cannot be revealed by biochemical protein-protein interaction analysis or genomic co-occurrence analyses alone. As shown in Fig. 4, depletion of a single CX3CR1⁺ host-cell population reprogrammed the magnitude of the majority of GIs, demonstrating that the epistatic fitness of tumor genotypes is conditional on the cellular composition of the metastatic niche (**Fig. 4**). Comparison of CX3CR1⁺- and MRC1⁺-cell depletion further demonstrated the causal influence of distinct metastatic niche components in rewiring the GI landscape, despite identical genetic perturbations in tumor cells (**Fig. 5**). Prior work has established differential epistasis under environmental and chemical perturbations (*16, 44*), context-dependent co-mutation across cancer types (*46*), in vivo combinatorial screening (*49, 50*), and human genetic-interaction maps generated in fixed cellular backgrounds (*51*). Our results extend these foundations by establishing host cell identity as a causal and anatomically addressable coordinate of genetic interaction. Niche conditioning resulted in a selective and highly structured GI landscape that cannot be explained by additive, independent gene effects. Among Pten partners, niche-dependent GIs varied in both magnitude and direction, directly linking the GI landscape to metastatic phenotype (**Figs. 4 and 5**). Thus, niche identity constitutes a third dimension of genetic interaction that determines the resulting GI landscape and, ultimately, metastatic outcomes.

Expanding Paget’s classic “seed and soil” model of metastasis, our temporal clonal dynamics and in vivo Perturb-seq analyses (**Fig. 3**) collectively suggest that the metastatic niche (the soil) imposes a two-layer evolutionary selection on disseminated tumor cells (the seeds). Clones with epistatic fitness advantages were selected during early colonization and remained stable during subsequent metastatic outgrowth. Yet, the dominant perturbation-associated transcriptional differences observed in vitro largely converged by the in vivo metastasis outgrowth stage, even though genotype-specific metastatic fitness had diverged during early colonization (**Fig. 3**).

Whereas metastatic adaptation is often framed in terms of transcriptional plasticity and dynamic cell-state transitions (*52, 53*), tumor cells that survived the earlier bottleneck of niche selection instead converged toward a narrower transcriptomic manifold, with only marginally detectable differences in inflammatory and interferon-response programs. Together, these results suggest a two-layer gate model in which the niche first selects among GI genotypes during early colonization and subsequently licenses or masks their epistatic phenotypic consequences during metastatic outgrowth. Importantly, this temporal uncoupling between early genotype-specific selection and later transcriptional convergence suggests that the cellular states observed in established metastases may incompletely capture the genotype-specific selective pressures that operated during earlier metastatic colonization.

Our host cell depletion experiments (the “backdrop” epistatic screens) further demonstrated how the same GI can be masked in one cellular environment and unmasked in another, providing a framework for interpreting differences in epistatic fitness across distinct host conditions. Most notably, MRC1⁺-cell depletion altered the Pten-Cdh1 GI and increased metastatic burden 14.6-fold (**Fig. 5**). This striking contrast argues against a generic tumor-suppressive effect of myeloid cells or an exclusively tumor-cell-autonomous effect of the Pten-Cdh1 interaction. Rather, it highlights specific control by MRC1⁺ cells, acting as the environmental “E,” over the phenotypic manifestation of G × G interactions. This G × G × E framework has implications beyond a specific gene pair, host cell type, or disease context. Therapy, infection, aging, and chronic inflammation can all alter host tissue composition and thereby change the epistatic fitness of existing tumor genotypes. Our findings therefore raise the possibility that perturbing the metastatic niche does more than alter its overall permissiveness for tumor growth. It may instead reorder the relative fitness of genetically distinct tumor clones. Consistent with the context-dependent and sometimes paradoxical consequences of therapeutically perturbing myeloid populations (*54–56*), our findings suggest that myeloid-targeted agents should not be assumed to exert uniform effects across tumor genotypes. Agents targeting myeloid populations, such as CSF1R or CCR2 inhibitors, could potentially reshape the GI landscape among genetically heterogeneous tumor cells, analogous to the effects observed in our genetic backdrop screens. Such remodeling could suppress the metastatic fitness of some genotypes while simultaneously exposing or enhancing latent fitness advantages in others. Given the extent to which host-cell composition rewired tumor-cell GIs in our study, it will be important to jointly evaluate co-occurring genetic alterations and niche composition, both before and after therapy, to predict genotype-specific therapeutic responses. Such studies could further reveal latent genetic dependencies exposed by treatment-induced remodeling of the GI landscape, thereby informing rational combination treatment strategies.

## Methods

### Ethical Regulations

All animal experiments were conducted according to IACUC protocols which were pre-approved by the UT Southwestern Medical Center IACUC committee (2022-103328) and the University of Notre Dame’s IACUC committee (18-05-468).

### Mice

Mice were housed with unlimited access to food and water in a temperature and humidity-controlled holding room (22 ± 2 °C, 50± 10% humidity) with a standard 12-hour light/dark cycle. Health checks were performed daily during in vivo experiments. Wild type C57BL/6J (B6) Cas12a central tolerance mice were 12-16 weeks old at the experiment onset. Control and depleted transgenic mice were age-matched, greater than 8 weeks of age. All experiments were carried out exclusively on female mice. B6.C-Tg(CMV-cre)1Cgn/J (CMV-Cre, 006054), B6.129P2(Cg)-Cx3cr1*^tm2.1(cre/ERT2)Litt^*/WganJ (Cx3cr1*^CreER^*knock-in/knock-out, 021160), and C57BL/6-Gt(ROSA)26Sor*^tm1(HBEGF)Awai/^*J (ROSA26iDTR, 007900) mice were purchased from Jackson Laboratory (Ben Harbor, ME). C57BL/6JSmo-Mrc1*^em2(2A-CreERT2)Smoc^* (Cat. NO. NMKI-220031) mice were purchased from Shanghai Model Organisms Center, Inc. (Shanghai, China). Cx3cr1*^CreER/+^*:R26*^iDTR/+^* and Mrc1*^CreER/+^*:R26*^iDTR/+^*double transgenic mice were produced in-house by breeding Cx3cr1*^CreER^*and Mrc1*^CreER^* with ROSA26*^iDTR^* mice. Cx3cr1*^CreER/+^* mice, used as controls in general myeloid depletion studies, were produced in-house by breeding Cx3cr1*^CreER^* mice to C57BL/6 mice. Mrc1*^CreER/+^* mice, used as controls in general lung interstitial macrophages depletion studies, were produced in-house by breeding Mrc1*^CreER^* mice to C57BL/6 mice. Specific numbers of mice used in each experiment are indicated in the corresponding figure legends.

### Generation of LSL-AsCas12a and constitutive AsCas12a mouse lines

LoxP-Stop-LoxP (LSL)-AsCas12a mice were generated by CRISPR/Cas9-mediated genome editing through pronuclear microinjection of C57BL/6J zygotes with a Rosa26-targeting crRNA/tracrRNA/Cas9 ribonucleoprotein complex and donor DNA, following established procedures. A Rosa26-targeting crRNA (5′-ACTCCAGTCTTTCTAGAAGA-3′) was used for targeted integration. The Rosa26-targeting donor vector was engineered to contain a Cre-dependent LSL expression cassette encoding AsCas12a-6×NLS-E174R/S542R together with an mCherry fluorescent reporter. The AsCas12a variant corresponds to the engineered AsCas12a-6×NLS-E174R/S542R sequence used in pRG232 (Addgene plasmid #149723).

Injected embryos were transferred to the oviducts of pseudopregnant CD-1 foster females using standard procedures. Founder mice were screened by PCR using primers targeting the AsCas12a transgene and the Rosa26 locus, and correct integration of the expression cassette was verified by Sanger sequencing across the targeted locus. Constitutive AsCas12a-expressing mice were generated by crossing LSL-AsCas12a mice with CMV-Cre mice (The Jackson Laboratory), thereby removing the LoxP-flanked transcriptional STOP cassette and enabling constitutive expression of AsCas12a and mCherry. These constitutive AsCas12a-expressing mice were used as Cas12a-tolerant hosts for in vivo CRISPR screening experiments, with the goal of minimizing host immune responses against the Cas12a protein carried by transplanted tumor cells.

Expression of the transgenic cassette was evaluated by fluorescence microscopy. mCherry fluorescence was examined in bone marrow-derived cells from constitutive AsCas12a-expressing mice and in whole-brain tissue from C57BL/6J, LSL-AsCas12a, and constitutive AsCas12a mice, providing cellular- and tissue-level validation of Cre-dependent transgene expression.

### Mouse primary brain astrocyte culture and validation of AsCas12a editing activity

Primary brain astrocyte cultures were established from constitutive AsCas12a-expressing mice to validate expression and genome-editing activity of the AsCas12a transgene at the cellular level. Brain tissue was dissected, mechanically dissociated and enzymatically digested, and the resulting cell suspension was filtered and maintained under astrocyte culture conditions. mCherry fluorescence was examined by fluorescence microscopy to confirm expression of the AsCas12a-associated reporter.

To assess the genome-editing activity of transgenic AsCas12a, primary astrocytes were transduced with lentiviral vectors expressing Cas12a crRNAs targeting Trp53, Pten, or Rb1. Following transduction, the corresponding genomic target regions were PCR-amplified using locus-specific primers (**table S7**) and subjected to amplicon sequencing. Editing efficiency and the distribution of insertion and deletion alleles at each target site were quantified using CRISPResso. Efficient on-target editing was observed across all three validation loci, with editing efficiencies of approximately 70–80%, confirming that the constitutively expressed AsCas12a transgene was functionally competent for crRNA-directed genome editing.

crTrp53 guide 1 + crPten guide 1 Top oligo

5’- **AGA T**CT TCC ACC CGG ATA AGA TGC TGG **TAA TTT CTA CTA TCG TAG AT**G CAG GTA GCT -3’

crTrp53 guide 1 + crPten guide 1 Bottom oligo

5’- TAT AGC TAC CTG C**AT CTA CGA TAG TAG AAA TTA** CCA GCA TCT TAT CCG GGT GGA AG -3’

crPten guide 1 + crRb1 guide 1 Top oligo

5’- ATA ATA ATA TAC A**AA ATT TCT CCT CTC GGA GAT** ACT TAC TGC TAT ACG TAG CCA TT -3’

crPten guide 1 + crRb1 guide 1 Bottom oligo

5’- **ATT C**AA TGG CTA CGT ATA GCA GTA AGT **ATC TCC GAG AGG AGA AAT TT**T GTA TAT TAT -3’

### Cell Lines

The TNBC E0771 cell line (CH3 Biosystems, Amherst, NY; Cat.No. 940001-Vial) was maintained in DMEM high glucose media supplemented with 10%FBS and 1% Penicilin/Streptomycin (Pen/Strep). HEK293T cells were cultured in DMEM high glucose supplemented with 10% FBS and 1% Pen/Strep. Cell cultures were grown at 37 °C, 5% CO_2_, and 95% humidity. All cell lines were tested and were negative for mycoplasma contamination using a routine mycoplasma PCR kit (ABM).

### Cas12a construct cloning

The AsCas12a-6×NLS-E174R/S542R coding sequence was PCR-amplified from pRG232 (Addgene plasmid #149723, a gift from Junwei Shi) using NEBNext Ultra II Q5 Master Mix (New England Biolabs, M0544) and cloned into the pENTR4-FLAG (w210-2) entry vector (Addgene plasmid #17423) using NEBuilder HiFi DNA Assembly Master Mix (New England Biolabs, E2621). The resulting AsCas12a entry construct was subsequently transferred into the doxycycline-inducible piggyBac destination vector PB-TAC-ERP2 (Addgene plasmid #80478) using Gateway LR Clonase II Enzyme Mix (Invitrogen, Thermo Fisher Scientific, 11791020), generating the final PB-232-mCherry construct. Correct plasmid assembly was confirmed by restriction enzyme digestion and Sanger sequencing.

### PiggyBac Transposon Transfection

To generate cells stably expressing inducible AsCas12a, target cells were seeded in 6-well plates 24 h before transfection. The PB-232-mCherry PiggyBac transposon plasmid encoding AsCas12a-6×NLS-E174R/S542R was co-transfected with the Super PiggyBac Transposase expression plasmid (System Biosciences, PB210PA-1) using CalFectin transfection reagent. The transfection mixture was added dropwise to the cells. At 12–16 h after transfection, the transfection medium was replaced with complete growth medium. At 48 h after transfection, cells were selected with 2 μg ml⁻¹ puromycin (Thermo Scientific Chemicals, J67236.XF) for at least 7 days to enrich for cells with stable integration of the PiggyBac transgene.

### Selection of candidate genes for the primary dual-crRNA screening library

Candidate genes for the primary combinatorial CRISPR screen were nominated based on genomic and transcriptomic alterations identified in clinical breast cancer metastases. At the genomic level, genes with a mutation frequency >15% across breast cancer metastases in the MSK-MET cohort, together with genes involved in recurrent co-occurring genetic events, were included as candidate genes. At the transcriptomic level, we analyzed the GSE110590 RNA-sequencing dataset comprising primary breast tumors and lung metastases from five patients (5 primary tumors and 8 lung metastases). Differential gene expression between primary tumors and lung metastases was analyzed using DESeq2. Genes with an adjusted P value <0.05 and log2 fold change <−1.5 in lung metastases relative to primary tumors were selected, yielding 228 candidate genes. Candidate genes nominated from the genomic and transcriptomic analyses were functionally annotated using Gene Ontology (GO) Biological Process terms and manually curated based on GO annotations, published literature, and established biological functions.

Genes were classified as tumor-intrinsic or niche-associated according to their predominant biological functions, with niche-associated genes encompassing immune, extracellular-matrix, stromal, and other microenvironment-related functions. Genes without sufficient functional evidence to support assignment to either category were excluded. The final gene set comprised 91 tumor-intrinsic genes and 78 niche-associated genes, which were used to construct the primary dual-crRNA screening library.

### Dual-crRNA library design and cloning

Two sequential dual-crRNA libraries were designed to balance screening breadth with in vivo library representation. The round-1 library was designed as a high-throughput, large-scale cross-program screen to broadly interrogate pairwise interactions between tumor-intrinsic and niche-associated genes. The library comprised 91 tumor-intrinsic genes at position 1 and 78 niche-associated genes at position 2, generating 7,098 designed cross-program gene combinations. In addition, 62 non-targeting control (NTC) crRNAs were included at each crRNA position, enabling gene-targeting crRNAs to be paired with NTCs for estimation of single-gene perturbation effects, as well as NTC–NTC combinations to provide non-targeting references.

Because metastatic colonization imposes a substantial population bottleneck, the large complexity of the round-1 library limited the representation that could be maintained for individual perturbations during in vivo selection. We therefore designed a 289-pair round-2 focused pairwise library based on genes and interactions prioritized from the round-1 screen. By substantially reducing library complexity, the round-2 screen enabled approximately 500-fold representation of individual perturbations in vivo and was used to validate and more deeply resolve the pairwise genetic interactions identified in the initial screen. The same 289-pair round-2 focused library was subsequently used in both host-depletion screens. Each depletion model was evaluated in an independent experiment comprising matched depleted and niche-intact wild-type host arms that received the same injected cell pool, thereby enabling comparison of genetic interactions under altered versus intact host-niche conditions.

Custom oligonucleotide pools encoding the dual-crRNA library were synthesized by Twist Bioscience. The oligonucleotide pool contained four classes of oligonucleotides corresponding to the forward and reverse strands of the first (position 1) and second (position 2) crRNA positions within the dual-crRNA cassette. Oligonucleotides were designed with BsmBI-compatible overhangs to facilitate combinatorial assembly of crRNAs at the two positions. The pooled oligonucleotides were phosphorylated using T4 Polynucleotide Kinase (T4 PNK; New England Biolabs, M0201S) and annealed to generate double-stranded dual-crRNA inserts containing random combinations of position 1 and position 2 crRNAs.

The annealed dual-crRNA inserts were cloned into the EFS-EGFP-P2A-Neo-WPRE-U6-AsCas12aDR (pRG212) backbone (Addgene plasmid #149722) using the NEBridge Golden Gate Assembly Kit (BsmBI-v2; New England Biolabs, E1602) at a 2:1 insert-to-vector molar ratio. Assembly reactions were incubated at 42 °C for 1 h followed by 60 °C for 5 min. The resulting plasmid library was transformed into Endura electrocompetent E. coli cells (Biosearch Technologies, 60242-1) and distributed across 20 LB agar plates containing ampicillin for overnight growth. Approximately 3 × 10^6^ -2 × 10^7^ colony-forming units (CFUs) were recovered per plate, yielding an estimated total of 6 × 10^7^ -4 × 10^8^CFUs and corresponding to approximately 240–1,600-fold representation of the ∼250,000 unique dual-crRNA library elements. Colonies were harvested by flushing the plates with prewarmed LB medium and subsequently expanded in liquid LB medium for 2 h. Plasmid DNA was purified using the ZymoPURE II Plasmid Maxiprep Kit (Zymo Research, D4202).

### Plasmid Library Amplification and Diversity Validation

Plasmid DNA was amplified using two step PCR to first amplify and then add indexing sequences to libraries. First round PCR was conducted using custom P5 and P7 adapters with overhangs compatible with Nextera indexing primers, followed by cleanup with Agencourt AMPure XP beads (Beckman Coulter, NC9933872) at a 0.9 × bead ratio to remove primers. Index PCR reaction was conducted as outlined in the Illumina 16S Metagenomic Sequencing Library Preparation manual (Illumina, Doc #15044223, page 11-12) using KAPA HiFi HotStart ReadyMix (Roche, KK2601) and Nextera XT Index kit v2 primers (Illumina, FC-131-1001). Final indexed libraries were quantified via the Qubit HS assay kit. Dual-crRNA library was run on TapeStation using the High Sensitivity D1000 assay (Agilent, 5067-5585, 5067-5594) and sequenced on an Illumina NovaSeq X Plus platform using 150-bp paired-end (PE150) reads. Demultiplexed fastq files were pre-processed using a customized script to estimate the percentage coverage based on the original crRNA whitelist.

### Lentivirus Production and Target Cell Transduction

pRG212-dual-crRNA-GFP and second-generation lentiviral packaging plasmids were transfected into HEK293T cells along with the CalFectin transfection reagent (SignaGen, SL100478). Lentivirus was harvested after 72 hours and concentrated using Lenti-X Concentrator (Takara Bio, 631231). Concentrated lentivirus was added to target cells along with 8 ug/mL polybrene. To achieve low MOI for CRISPR screening experiments, E0771 cancer cells were infected at a low multiplicity of infection (MOI) of 0.15, as calculated by the percentage of GFP-positive cells 48 hours post transduction. 48 hours post transfection, cells were selected with G418 (Sigma-Aldrich, 4727878001) for 7 days.

### Establishment and Monitoring of Experimental Metastasis

For the LungMets model, 1x10^6^ E0771.mCherry cells were injected intravenously via the retro-orbital sinus to ensure adequate library representation (>500X coverage) during the second-round CRISPR screening experiments. Mice were sacrificed 3 days and 18 days after injection. For experimental metastasis studies, maximal permitted burden was defined by humane endpoint criteria defined by the IACUC institutional committee.

### Genomic DNA (gDNA) Amplicon Library Preparation

Genomic DNA was extracted from cultured cells and digested tissues using either the Quick-DNA Midiprep Kit (Zymo Research, D4075) or the Quick-DNA Miniprep Plus Kit (Zymo Research, D4068), depending on the starting cell number. Extracted DNA yield was quantified via a NanoDrop spectrophotometer, and samples were normalized to the same, lowest input concentration prior to PCR. Two rounds of PCR amplification were performed on > 200 ng starting gDNA material. The first round was utilized for amplicon amplification with nested primers for 15-20 cycles using a high-fidelity hot start polymerase, followed by purification with a 0.9x bead-to-sample AMPure XP bead ratio. Eluted amplicon DNA was quantified using the Qubit high sensitivity (HS) assay kit (Thermo Fisher Scientific, Q33231) and concentrations normalized prior to indexing. Approximately 5 ng of DNA was carried into the index PCR rection, which was conducted similarly to the plasmid library construction protocol described above. Index products were purified using the same AMPure XP beads ratio to remove excess primers. Library quantity was assessed via the Qubit HS assay kit and analyzed on a TapeStation instrument using the High Sensitivity D1000 assay to confirm expected library size and distribution. Dual-crRNA library was sequenced on a NovaSeq-X instrument. High quality reads were filtered and processed using the MAGeCK pipeline to identify enriched or depleted crRNAs. Primers used for amplification were as follows:

Round 1 PCR REV, Tm: 65.0:

5’ – GTC TCG TGG GCT CGG AGA TGT GTA TAA GAG ACA GGT CTC AAG ATC TAG TTA CGC CAA G– 3’

Round 1 PCR FWD, Tm: 65.0:

5’ – ACG ACG CTC TTC CGA TCT GAC GAA ACA CCG AAT TTC TAC TCT– 3’

### Combinatorial CRISPR screen data processing and genetic interaction analysis

FASTQ files generated from the combinatorial CRISPR screens were processed using MAGeCK (v0.5.6). Dual-crRNA construct abundance was quantified using mageck count with total-count normalization (--norm-method total) against the custom dual-crRNA library. A pseudocount of 1 was added to zero counts. Differential representation of individual dual-crRNA constructs between experimental samples and the injected cell pool was subsequently assessed using mageck test, with non-targeting constructs specified as normalization controls (--norm-method control) and multiple-testing correction performed using the false discovery rate (FDR) method (--adjust-method fdr).

To ensure adequate baseline representation, constructs were required to have at least five reads in the injected cell pool, and gene pairs were retained when at least five independent constructs met this criterion. No minimum-read threshold was imposed on lung or in vitro samples, allowing constructs that became strongly depleted or undetectable during selection to remain represented in downstream analyses. Following MAGeCK analysis, informative dual-crRNA constructs were selected based on an absolute log2 fold change (LFC) >0.5, either a lower-tail P value (p.low) <0.05 or an upper-tail P value (p.high) <0.05, and FDR <0.25. This filtering yielded 985 dual-crRNA constructs for downstream gene-pair analysis.

Multiple dual-crRNA constructs targeting the same gene pair were collapsed to the gene-pair level by inverse-variance-weighted aggregation of guide-level LFCs, retaining at most the 25 highest-abundance guides per pair. Single-knockout effects were estimated from constructs pairing a gene-targeting crRNA with a non-targeting crRNA. Genetic interaction (GI) scores were calculated as the deviation of the observed double-knockout effect from the additive expectation of the corresponding single-knockout effects:

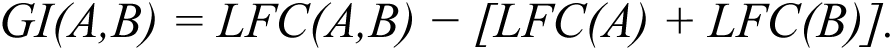

Positive and negative GI scores indicate deviations above and below the additive expectation, respectively. GI standard errors were propagated from the double-knockout and corresponding single-knockout estimates. Two-sided P values were calculated from the resulting z scores and adjusted for multiple testing using the Benjamini–Hochberg procedure.

### Individual Gene pair Knockout Guide Cloning

The pRG212 plasmid (Addgene # 149722) was first modified to insert two BsmBI recognition sites between the AS_DR and DR_10 sequences through a NEBuilder HiFi DNA Assembly (NEB, E5520S) reaction. Then, CRISPR RNA (crRNA) sequences were designed using CRISPick (Mouse GRCm38 reference genome), oligos synthesized by IDT with BsmBI overhangs, annealed, and cloned into the modified pRG212 plasmid using the NEBridge Golden Gate Assembly Kit (NEB, E1602S). The resulting plasmids were first transformed into competent *E. coli*, plated on LB-ampicilin (AMP) plates, and grown at 30 °C, before plasmid extraction. Below are the crRNA sequences used for gene targeting, including the necessary BsmBI overhangs:

crPten guide 1 Top oligo

5’-**AGA T**GC AGG TAG CTA TAA TAA TAT ACA **AAA TTT CTC CTC TCG G** -3’

crPten guide 1 Bottom oligo

5’-**ATT T**TG TAT ATT ATT ATA GCT ACC TGC -3’

crPten guide 4 Top oligo

5’-**AGA T**GC TGG CAG ACC ACA AAC TGA GGA **AAA TTT CTC CTC TCG G** -3’

crPten guide 4 Bottom oligo

5’-**ATT T**TC CTC AGT TTG TGG TCT GCC AGC -3’

crChek2 guide 2 Top oligo

5’-**AGA T**CC TTT CCC AAT AAG CTC GGT ATT **AAA TTT CTC CTC TCG G** -3’

crChek2 guide 2 Bottom oligo

5’-**ATT T**AA TAC CGA GCT TAT TGG GAA AGG -3’

crChek2 guide 3 Top oligo

5’-**AGA T**AA CGT TAC TAC TTA CAG TTC CAG **AAA TTT CTC CTC TCG G** -3’

crChek2 guide 3 Bottom oligo

5’-**ATT T**CT GGA ACT GTA AGT AGT AAC GTT -3’

crCdh1 guide 1 Top oligo

5’-**AGA T**GT GTC TAC AGA GTT ACC CTA CAT **AAA TTT CTC CTC TCG G** -3’

crCdh1 guide 1 Bottom oligo

5’-**ATT T**AT GTA GGG TAA CTC TGT AGA CAC -3’

crCdh1 guide 3 Top oligo

5’-**AGA T**AT GAC ACG GCA TGA GAA TAG AGC **AAA TTT CTC CTC TCG G** -3’

crCdh1 guide 3 Bottom oligo

5’-**ATT T**GC TCT ATT CTC ATG CCG TGT CAT -3’

crCx3cl1 guide 3 Top oligo

5’-**AGA T**GT GAG GGC AGC AGC CTG GTG ATC -3’

crCx3cl1 guide 3 Bottom oligo

5’-**ATT C**GA TCA CCA GGC TGC TGC CCT CAC **ATC TCC GAG AGG AGA A** -3’

crCx3cl1 guide 5 Top oligo

5’-**AGA T**CT CTG GGG TTG AAG GTG AAG TAG -3’

crCx3cl1 guide 5 Bottom oligo

5’-**ATT C**CT ACT TCA CCT TCA ACC CCA GAG **ATC TCC GAG AGG AGA A** -3’

crCxcr5 guide 2 Top oligo

5’-**AGA T**CT GGG AGA AGG TGC ACT GTG GTA -3’

crCxcr5 guide 2 Bottom oligo

5’-**ATT C**TA CCA CAG TGC ACC TTC TCC CAG **ATC TCC GAG AGG AGA A** -3’

crCxcr5 guide 4 Top oligo

5’-**AGA T**CT CTA CCA CAT CGG GGG CTT CCT -3’

crCxcr5 guide 4 Bottom oligo

5’-**ATT C**AG GAA GCC CCC GAT GTG GTA GAG **ATC TCC GAG AGG AGA A** -3’

crTlr7 guide 2 Top oligo

5’-**AGA T**GT GAG AGA ACC TTC AAA TTT CTC -3’

crTlr7 guide 2 Bottom oligo

5’-**ATT C**GA GAA ATT TGA AGG TTC TCT CAC **ATC TCC GAG AGG AGA A** -3’

crTlr7 guide 5 Top oligo

5’-**AGA T**AG TCA GAG AGT CCA CTA AAG CTT -3’

crTlr7 guide 5 Bottom oligo

5’-**ATT C**AA GCT TTA GTG GAC TCT CTG ACT **ATC TCC GAG AGG AGA A** -3’

crNTC-6 Top oligo

5’-**AGA T**GA CCG TTG TAT CGC GAT CAA -3’

crNTC-6 Bottom oligo

5’-**ATT C**TT GAT CGC GAT ACA ACG GTC **ATC TCC GAG AGG AGA A** -3’

crNTC-17 Top oligo

5’-**AGA T**AC GCG CTA TTA TCG TAG GTG -3’

crNTC-17 Bottom oligo

5’-**ATT C**CA CCT ACG ATA ATA GCG CGT **ATC TCC GAG AGG AGA A** -3’

crNTC-30 Top oligo

5’-**AGA T**TG GCG CGT AAT CGC TAT TCG **AAA TTT CTC CTC TCG G** -3’

crNTC-30 Bottom oligo

5’-**ATT T**CG AAT AGC GAT TAC GCG CCA -3’

crNTC-43 Top oligo

5’-**AGA T**TC TCG TAT CGC GAT TGT TAA **AAA TTT CTC CTC TCG G** -3’

crNTC-43 Bottom oligo

5’-**ATT T**TT AAC AAT CGC GAT ACG AGA -3’

### Validation of editing efficiency in individual knockout cell lines

Editing efficiency of individual dual-crRNA constructs used for downstream functional validation was assessed by targeted amplicon sequencing. Genomic regions spanning the corresponding Cas12a target sites were PCR-amplified from edited E0771 cells using locus-specific primers (**table S7**), and editing outcomes were quantified using CRISPResso. The percentage of modified reads and the distribution of insertion and deletion alleles were determined independently for each targeted locus. For dual-knockout constructs, editing efficiency was evaluated at both target sites.

### Cell Preparation for Perturb-Seq

Tumor cells for in vivo and matched in vitro Perturb-seq were prepared in parallel using the same focused perturbation pools. For the in vivo arm, tumor-bearing mice were euthanized by cervical dislocation 14 days after tumor cell injection. Mice were perfused with 5 ml of ice-cold 1× PBS, and lungs were collected and examined for GFP fluorescence using a fluorescence dissecting microscope. Lungs with detectable GFP-positive tumor burden were minced into small pieces and enzymatically dissociated in serum-free RPMI 1640 containing 1× collagenase/hyaluronidase (STEMCELL Technologies, 07912) and 0.15 mg ml⁻¹ DNase I (Roche, MilliporeSigma, 10104159001). Digested tissues were passed through a 70-μm cell strainer, washed with serum-free RPMI 1640, and treated with 1× RBC Lysis Buffer (BioLegend, 420301) to remove erythrocytes. Cells were subsequently stained with Zombie Aqua Fixable Viability Kit (BioLegend, 423101) for exclusion of dead cells.

For the matched in vitro arm, aliquots of the corresponding perturbation pools were maintained in culture in parallel with the in vivo experiment for the same duration. On day 14, coincident with collection of the lung metastasis samples, cultured cells were detached by trypsinization, collected, and washed to generate single-cell suspensions. The in vitro samples were then processed in parallel with the in vivo samples for cell hashing and fluorescence-activated cell sorting. Thus, the in vitro and in vivo Perturb-seq samples were collected at the same experimental endpoint and subjected to the same downstream sample-multiplexing and sorting workflow. This paired design enabled direct comparison of perturbation-associated transcriptional states in culture and following metastatic colonization in vivo.

For sample multiplexing, both in vivo- and in vitro-derived cell suspensions were labeled with TotalSeq-C Mouse Hashtag antibodies (BioLegend) according to the CITE-seq/cell-hashing workflow. Briefly, cells were incubated with FcR Blocking Reagent (Miltenyi Biotec, 130-092-575) in 50 μl of cell staining buffer for 20 min on ice, followed by staining with 1.0 μg per test of the indicated TotalSeq-C hashing antibody for 25 min on ice. The hashing antibodies including: M-HTO-1 (M1/42; 30-F11, 155861), M-HTO-2 (M1/42; 30-F11, 155863), M-HTO-3 (M1/42; 30-F11, 155865), M-HTO-4 (M1/42; 30-F11, 155867), M-HTO-9 (M1/42; 30-F11, 18446), M-HTO-12 (M1/42; 30-F11, 22005),

Following staining, cells were washed three times with cell staining buffer containing 2 mM EDTA, 1 mM EDTA, and no EDTA, respectively. Viable GFP-positive tumor cells (Zombie Aqua⁻GFP⁺) were isolated using a BD FACSAria cell sorter and subsequently prepared for loading onto a 10x Genomics Chromium Controller.

### Perturb-seq Library Preparation and Illumina Sequencing

Perturb-seq libraries were prepared using the Chromium GEM-X Single Cell 5′ Reagent Kits v3 with Feature Barcode technology for CRISPR Screening and Cell Surface Protein (10x Genomics), with modifications for the capture and amplification of AsCas12a crRNAs. The Chromium GEM-X Single Cell 5′ Kit v3 (PN-1000699), Chromium GEM-X Single Cell 5′ Chip Kit (PN-1000698), Chromium GEM-X Single Cell 5′ Feature Barcode Kit v3 (PN-1000703), and Dual Index Kit TT Set A (PN-1000215) were used for GEM generation, barcoding, cDNA amplification, and library construction. The workflow was used to generate 5′ gene expression, CRISPR screening, and cell surface protein Feature Barcode libraries.

To enable capture of AsCas12a crRNAs, a custom CS1_enrichment primer (5′-GTGACTGGAGTTCAGACGTGTGCTCTTCCGATCTTTGCTAGGACCGGCCTTAAAGC-3′) was introduced during GEM reverse transcription (GEM-RT). Specifically, 0.5 μl of the CS1_enrichment primer (10 nM) was spiked into the Poly-dT RT Primer B (10x Genomics, PN-2001110) during GEM-RT.

For CRISPR screening library construction, the Feature PCR step was modified to specifically amplify the captured AsCas12a crRNA-derived cDNA. Feature PCR was performed using the standard Read1T primer from the 10x Genomics workflow (5′-ACACTCTTTCCCTACACGACGC-3′) together with the custom CS1_enrichment primer.

Primer volumes and concentrations were maintained as specified for the corresponding primers in the manufacturer’s protocol. Subsequent CRISPR library cleanup, sample indexing, size selection, and quality control were performed according to the manufacturer’s protocol.

For sample multiplexing, six TotalSeq-C Mouse Hashtag Oligonucleotide (HTO)-labeled samples were processed to generate Cell Surface Protein Feature Barcode libraries. HTO libraries were prepared without modification according to the manufacturer’s Cell Surface Protein library construction protocol, including sample index PCR, post-PCR size selection, and library quality control.

The quality and quantity of the three individual libraries (transcriptome, HTO, and CRISPR) were assessed via Qubit and TapeStation prior to sequencing. Libraries were pooled for sequencing on an Illumina NovaSeq X Plus platform with paired end reads (PE150).

### Perturb-seq General Data Analysis

Raw FASTQ files generated from the sequencing runs, including gene expression (mRNA), hashtag oligonucleotide (HTO), and CRISPR guide capture libraries, were processed using Cell Ranger (v8.0.1; 10x Genomics) to generate feature-barcode count matrices. Gene expression reads were aligned to a custom mouse reference genome supplemented with sequences corresponding to the fluorescent reporter transcripts used in this study. The resulting Gene Expression, Antibody Capture, and CRISPR Guide Capture count matrices were imported into R and analyzed using Seurat (v5.0.1).

For sample demultiplexing, HTO counts were stored as a separate Seurat assay and normalized using centered log-ratio (CLR) transformation. HTO identities were assigned using HTODemux with a positive quantile of 0.9. Cells classified as HTO doublets or HTO-negative were excluded from subsequent analyses.

For perturbation assignment, CRISPR Guide Capture counts were stored as a separate Seurat assay and normalized using centered log-ratio (CLR) transformation. Because the two crRNAs occupying positions 1 and 2 of each dual-crRNA construct were captured and sequenced together within the same read, each detected dual-crRNA sequence was treated as a single perturbation identity without requiring computational pairing of independently detected crRNAs. Initial classification of CRISPR Guide Capture signals was performed using MULTIseqDemux. A cell was assigned to a dual-crRNA construct when it contained at least three total guide-capture counts in the CRISPR Guide Capture count matrix and a single dual-crRNA identity accounted for >75% of the total guide-capture counts detected in that cell. When multiple dual-crRNA identities were detected, the most abundant identity was assigned if it exceeded the 75% threshold. Cells with fewer than three total guide-capture counts or without a dominant dual-crRNA identity exceeding the 75% threshold were classified as CRISPR-negative and excluded from downstream transcriptomic analyses. Individual dual-crRNA constructs containing different crRNA sequences targeting the same gene or gene pair were subsequently collapsed into gene-level perturbation groups for downstream analyses. The resulting perturbation annotations were incorporated into the Seurat metadata prior to gene expression analysis.

For gene expression quality control, the percentage of reads mapping to mitochondrial genes was calculated for each cell. Cells expressing fewer than 200 or more than 7,500 genes, or with >5% mitochondrial reads, were excluded. Gene expression data were normalized using SCTransform with vst.flavor = “v2”, using 3,000 variable features and regressing out the percentage of mitochondrial reads. The glmGamPoi implementation was used when available.

Data generated from the two independent Chromium 10x lanes, corresponding to the synergistic and buffering perturbation panels, were integrated using an SCTransform-based integration workflow. Principal component analysis (PCA) was performed on the integrated dataset for dimensionality reduction, followed by construction of a shared nearest-neighbor graph, unsupervised clustering, and uniform manifold approximation and projection (UMAP) for visualization.

### In Vivo Myeloid Depletion

To deplete the general Cx3cr1+ myeloid population in Cx3cr1*^CreER/+^*:R26*^iDTR/+^*mice and lung interstitial macrophage population in Mrc1*^CreER/+^*:R26*^iDTR/+^*mice, mice were first administered 75mg kg-1 Tamoxifen (Sigma, T5648) dissolved in corn oil (Sigma, C8267) by intraperitoneal injection once daily for 7 consecutive days before tumor cell inoculation, and then daily thereafter, to induce genetic recombination. Mice were administered 1 μg Diphtheria Toxin (Sigma, D0564) dissolved in 100 μL sterile saline by intraperitoneal injection three consecutive days prior to tumor cell injection and every other day thereafter. Tamoxifen and Diphtheria Toxin doses were reduced or skipped in some cases when mice were deemed too unhealthy and unable to handle further drug administration.

### Flow Cytometry

Samples were extracted and pre-processed similarly as the Perturb-seq samples. Following tissue digestion of lung tissues and RBC lysis, digested samples were washed in 1 × Flow Staining Buffer (Tonbo Biosciences, TNB-4222-L500). For viability assessment, cells were incubated with ZombieAqua Fixable Viability Kit (BioLegend, 423102) at a 1:750 dilution in 1 × PBS for 20 minutes at room temperature. Following viability staining, cells were blocked with 1 × FcR Blocking Reagent (Miltenyi Biotec, 130-092-575) to minimize nonspecific antibody staining.

Cells were stained with fluorochrome-conjugated surface antibodies at manufacturer-recommended concentrations. Depending on the experimental panel, antibodies included PerCP/Cyanine5.5 anti-mouse CD45 (BioLegend, 103131, 30-F11), Brilliant Violet 510™ anti-mouse CD3 (BioLegend, 100233, 17A2), Alexa Fluor® 700 anti-mouse CD11b (BioLegend, 101222, M1/70), APC/Cyanine7 anti-mouse CD4 (BioLegend, 100525, GK1.5), PE/Cyanine7 anti-mouse CD8a (BioLegend, 162311, 53-6.7), Brilliant Violet 421™ anti-mouse F4/80 (BioLegend, 123131, BM8), Brilliant Violet 605™ anti-mouse CD11c (BioLegend, 117333, N418), PE anti-mouse Ly6C (BioLegend, 128007, HK1.4), PE/Dazzle™ 594 anti-mouse NK-1.1 (BioLegend, 108747, PK136), APC anti-mouse I-A/I-E (MHC class II) (BioLegend, 107613, M5/114.15.2), and Alexa Fluor® 488 anti-mouse CX3CR1 (BioLegend, 149021, SA011F11).

For macrophage profiling, antibodies included PerCP/Cyanine5.5 anti-mouse CD45 (BioLegend, 103131, 30-F11), PE/Cyanine7 anti-mouse Ly-6G (BioLegend, 127617, 1A8), Alexa Fluor® 488 anti-mouse/human CD11b (BioLegend, 101219, M1/70), Brilliant Violet 421™ anti-mouse CD11c (BioLegend, 117329, N418), APC/Cyanine7 anti-mouse I-A/I-E (MHC class II) (BioLegend, 107627, M5/114.15.2), PE anti-mouse CD64 (FcγRI) (BioLegend, 139303, X54-5/7.1), Brilliant Violet 650™ anti-mouse CD24 (BioLegend, 101857, M1/69), PE/Dazzle™ 594 anti-mouse CD170 (Siglec-F) (BioLegend, 155529, S17007L), and Alexa Fluor® 647 anti-mouse CD206 (MMR) (BioLegend, 141711, C068C2). Cells were then washed with staining buffer, followed by analysis on a Cytek Northern Lights flow cytometer. FCS files were exported and analyzed using FlowJo (BD Biosciences).

### Estrogen-response proliferation assay

E0771 cells carrying four CRISPR knockout genotypes, including the double non-targeting control (NT_NT), Chek2 single knockout (Chek2_NT), Cx3cl1 single knockout (NT_Cx3cl1), and Chek2/Cx3cl1 double knockout (Chek2_Cx3cl1), were used to assess genotype-dependent proliferative responses to estrogen signaling. Cells were maintained in the complete Dulbecco’s Modified Eagle’s Medium (DMEM) supplemented with 10% fetal bovine serum (FBS) and 1% penicillin-streptomycin at 37°C in a humidified incubator with 5% CO₂.

Prior to hormone treatment, cells were switched to Minimum Essential Medium (MEM) phenol red-free medium supplemented with 10% charcoal-stripped FBS for 48 hours to minimize background estrogenic activity. Cells were then seeded into 96-well plates at 2000 cells per well in 200 μL medium per well and allowed to adhere for 24h before treatment. Where applicable, cells were maintained in hormone-depleted medium throughout the treatment period.

Cells of each genotype were subjected to five experimental conditions: baseline (no treatment), vehicle, fulvestrant (10 nM; MedChemExpress; HY-13636), 17β-estradiol (E2, 10 nM; Sigma-Aldrich; E2758), or combined E2 and fulvestrant (10 nM each). Stock solutions of E2 and fulvestrant were prepared in EtOH and diluted in culture medium immediately before use. Vehicle-treated cells received an equivalent final concentration of the corresponding solvent. Cells were exposed to the indicated treatments for 2 days.

Following treatment, cell proliferation was quantified using direct cell counting. The percent proliferation change was calculated as:

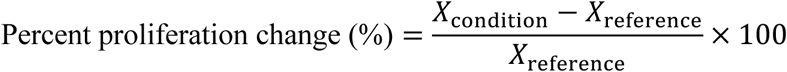

Each genotype and treatment condition was assessed in three independent biological replicates (n = 3). Data are presented as the percent proliferation change, with box plots indicating the interquartile range and individual points representing biological replicates.

### Immunofluorescence staining of lung metastatic lesions

Lung tissues containing metastatic lesions derived from GFP-labeled E0771 Pten single-knockout (Pten KO) or Pten/Cdh1 double-knockout (Pten+Cdh1 DKO) cells were collected 3 days after retro-orbital injection. Mice were euthanized and perfused with 10 mL ice-cold phosphate-buffered saline (PBS). Lungs were harvested and fixed in 4% paraformaldehyde (PFA) in PBS for 18 hours at 4°C. Fixed tissues were sequentially cryoprotected in 10%, 20%, and 30% sucrose in PBS overnight, respectively, at 4°C. Tissues were subsequently embedded in optimal cutting temperature (OCT) compound (Fisher HealthCare, 4585), frozen, and sectioned at 8μm thickness using a cryostat (Leica CM1950). Sections were mounted onto Superfrost Plus™ Microscope Slides (fisherbrand; 1255015) and stored at -20°C until staining.

For immunofluorescence staining, cryosections were air-dried for 10 min, washed with PBS, and permeabilized in 1% normal serum with 0.2% Triton X-100 in PBS for 10 min. Sections were blocked in 5% normal serum with 0.2% Triton X-100 in PBS for 1h at room temperature and subsequently incubated with primary antibodies against Ki67 (CST, 9129S; dilution 1:500) and MRC1/CD206 (R&D systems, AF2535; 5µg/mL) in 1% normal serum with 0.1% Triton X-100 in PBS (PBST) overnight at 4°C. Sections were then washed three times for 5min each with 0.1% PBST and incubated with the appropriate species-specific fluorophore-conjugated secondary antibodies (Alexa Fluor® 647 AffiniPure® Donkey Anti-Rabbit IgG (H+L), Jackson ImmunoResearch, 711-605-152; dilution 1:600 and Donkey anti-Goat IgG (H+L) Cross-Adsorbed Secondary Antibody Alexa Fluor™ 555, Invitrogen; 3µg/mL) for 1.5h at room temperature protected from light. Sections were then washed three times for 5min each with 0.1% PBST. Nuclei were counterstained with DAPI (Invitrogen, D1306; 1µg/mL) for 5min. Sections were washed three times for 5min each with PBS and mounted using SlowFade™ Diamond Antifade Mountant (Invitrogen, S36972).

Metastatic lesions were identified based on GFP fluorescence from the labeled E0771 tumor cells together with tissue morphology. Fluorescence images were acquired using a Nikon ECLIPSE Ti2 equipped with a 40× objective. Corresponding fluorescence channels were acquired using identical imaging settings across experimental groups.

### Clinical genomic and transcriptomic data analysis

#### Clinical cohorts and genomic alteration analysis

Publicly available genomic datasets were obtained from The Cancer Genome Atlas (TCGA) PanCancer Atlas and the MSK-MET metastatic cancer cohort through cBioPortal. Pan-cancer analyses included ten solid-tumor lineages (BRCA, LUAD, LUSC, COADREAD, STAD, HNSC, BLCA, PRAD, OV, and LIHC) using harmonized MC3 somatic mutation calls. Pairwise somatic co-occurrence was evaluated across a curated 53-gene cancer-driver panel using somaticInteractions in the R package maftools, which performs pairwise Fisher’s exact tests followed by Benjamini–Hochberg (BH) correction. Gene pairs with adjusted P < 0.05 in the pooled pan-cancer analysis were retained, and recurrence was defined as independent detection at adjusted P < 0.10 within individual tumor lineages.

#### Breast cancer-specific genomic analyses used the MSK-MET breast cancer cohort

The distant-metastatic cohort comprised 1,512 cases with documented involvement of bone, liver, lung, or brain. Pairwise analyses were performed using a predefined panel of 20 canonical breast cancer driver genes representing distinct genomic loci. A gene was considered altered when a tumor harbored either a non-silent somatic mutation or a high-level copy-number alteration (amplification or deep deletion); shallow copy-number changes were excluded. Pairwise co-occurrence and mutual exclusivity were assessed across all 190 gene pairs using two-sided Fisher’s exact tests followed by BH correction, with significant associations defined at FDR < 0.10. Pairwise effect sizes were represented by log odds ratios with a Haldane correction of 0.5 applied to each cell of the 2 × 2 contingency table.

#### Metastatic site-specific genomic analysis

Metastatic site was defined by the anatomical origin of the sequenced metastatic specimen rather than the patient’s overall history of metastatic involvement. Lung-, liver-, bone-, and brain-derived metastatic specimens were analyzed separately. For each site, Haldane-corrected pairwise log odds ratios were calculated across the 20-gene driver panel, and divergence in co-occurrence structure between the indicated metastatic site and other metastatic specimens was quantified as the mean absolute difference in log odds ratio across all 190 gene pairs. Statistical significance was assessed using one-sided label permutations. To account for differences in cohort composition, matched permutations were performed within strata defined by metastatic burden and intrinsic breast cancer subtype. A total of 2,500 permutations were performed for each metastatic site and covariate model, and BH correction was applied across the four metastatic sites within each model.

#### Primary breast tumor and lung metastasis transcriptomic analysis

Transcriptomic analyses were performed using the GSE110590 breast cancer RNA-sequencing dataset. The paired subset comprised five patients with matched primary breast tumors and lung metastases, yielding 5 primary tumors and 8 lung metastatic lesions. Unsupervised principal-component analysis was performed on the variance-filtered expression matrix after gene-wise scaling. Differential expression between primary tumors and lung metastases was evaluated using limma, with within-patient correlation estimated using duplicateCorrelation and patient identity specified as the blocking factor. Moderated statistics were calculated using eBayes(trend = TRUE), and differentially expressed genes were defined by BH-adjusted P < 0.05 and |log2 fold change| >1. Significant genes were subsequently assigned to curated Hallmark functional modules.

To examine pathway-level transcriptional rewiring, per-sample activity scores for MSigDB Hallmark gene sets were calculated as the mean of gene-wise standardized expression values among detected genes in each module. Modules containing at least 10 detected genes were retained. Spearman correlation matrices of module activity scores were calculated separately for primary tumors and lung metastases, and differential module co-activation was quantified as Δρ = ρ_lung metastasis − ρ_primary for each module pair. This analysis was used descriptively to identify pathway relationships altered during lung metastasis.

#### DawnRank-based gene co-activation analysis

DawnRank scores were obtained from Supplementary Table 5 of GSE110590 and used without additional normalization. An author-curated panel of 35 tumor-intrinsic and niche-associated genes was intersected with the available DawnRank scores, retaining 27 genes (10 tumor-intrinsic and 17 niche-associated). Pairwise Pearson correlations were calculated across 8 lung metastatic lesions from 5 patients. Correlations with |r| ≥ 0.55 were highlighted for visualization; this threshold was used as a descriptive display criterion rather than a statistical significance cutoff.

### Statistics & Reproducibility

Sample sizes were not predetermined using formal statistical methods and were selected based on experimental feasibility and established practices for the corresponding assays. The number of independent biological replicates and the definition of n are indicated in the corresponding figure legends. For animal experiments, individual mice were considered biological replicates unless otherwise stated. Statistical analyses were performed using GraphPad Prism and R, with specific computational methods and software packages described in the corresponding Methods sections. Comparisons between two independent experimental groups were performed using two-sided Student’s t-tests or Welch’s t-tests, as appropriate. For analyses involving multiple hypotheses, P values were adjusted for multiple testing using the Benjamini–Hochberg procedure unless otherwise specified. Differential gene-expression analyses were performed using the statistical frameworks implemented in DESeq2, limma, or the indicated single-cell analysis workflow, as appropriate for each dataset. Pairwise genomic co-occurrence analyses were evaluated using Fisher’s exact tests, and permutation-based analyses were used where indicated. Combinatorial CRISPR-screen statistics were derived using MAGeCK and the genetic-interaction framework described above. Exact statistical tests, significance thresholds, sample sizes, and definitions of error bars are provided in the corresponding figure legends and Methods sections. All reported replicates represent biologically independent samples unless explicitly stated otherwise.

## Acknowledgments

We thank the following core facilities for their support: the Simmons Comprehensive Cancer Center, the UT Southwestern Animal Resource Center, the UT Southwestern Flow Cytometry Core, the Children’s Research Institute Mouse Genome Engineering Core, the UT Southwestern Texas Cancer Cell Imaging Core, the University of Notre Dame Genomics and Bioinformatics Core Facility, and the Indiana University School of Medicine Center for Medical Genomics. The computational component of this project was supported in part by the BioHPC high-performance computing facility at UT Southwestern.

## Author Contributions Statement

Conceptualization and experiment design, X.L., L.H., and S.Z.; Data analysis, X.L., L.H., D.G., J.L., L.X., and S.Z.; Experiments and methodology, X.L., L.H., H.Z., M.Z., S.M.G., W.M., X.Y., E.A., H.M.B., K.M.D., L.W., G.C.H., and S.Z; Manuscript writing and revision, X.L., L.H., and S.Z.; Study supervision, S.Z. We attest that all coauthors reviewed and pre-approved the manuscript before submission.

## Data, code, and materials availability

The source data underlying all figures are provided with the paper. Perturb-seq datasets containing CRISPR guide and HTO multiplexing information have been deposited in the Gene Expression Omnibus under accession nos. GSE322808 and GSE322809. The processed single-cell objects derived from them are deposited at Zenodo (doi:10.5281/zenodo.21384699).

Construct-level and gene-level count matrices for the combinatorial CRISPR screens, from which all reported genetic-interaction and enrichment statistics are computed, are provided in the analysis repository. Analysis and plotting code for all figures, together with the package versions used, is available at https://github.com/S-Zhang-Lab/Epistasis-manuscript-code under the MIT license and archived at Zenodo. Clinical genomic data analyzed here are publicly available: the TCGA PanCancer Atlas and MSK-MET (2021) cohorts through cBioPortal and paired primary and metastatic transcriptomes under GEO accession GSE110590. Mouse lines, cell lines, and the combinatorial crRNA libraries generated in this study are available from the corresponding author under a standard materials transfer agreement.

## Supplementary Materials

Figs. S1 to S5

Tables S1 to S7

