## Supplemental Figures for "Niche-dependent epistasis within the cancer genome shapes metastatic fitness"

### Supplementary Fig. 1

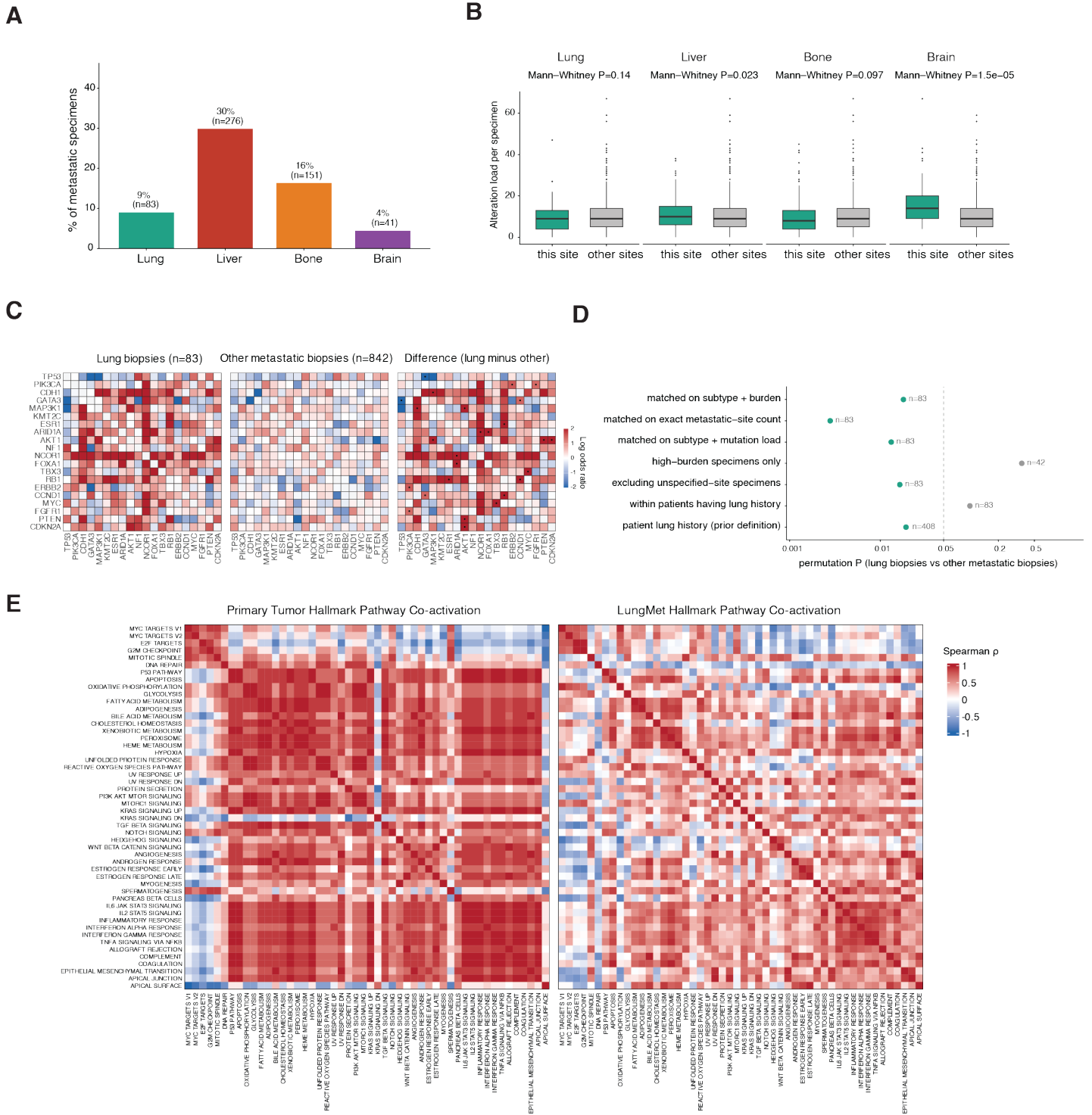

**Fig. S1. Cohort composition and robustness of the lung co-occurrence divergence (related to Fig. 1).** MSK-MET breast, metastasis-derived specimens ($n = 925$) analyzed using the biopsy site of the specimen, not the patient's recorded history (only 83 of 408 lung-history specimens, 20%, are lung biopsies). Co-occurrence is the Haldane-corrected pairwise log odds ratio over 20 drivers. Divergence is the mean absolute difference across the 190 gene pairs. **(A)** Biopsy-site composition (lung 83, liver 276, bone 151, brain 41). **(B)** Per-specimen alteration load (nonsilent mutations plus high-level copy-number alterations on the targeted panel, not a per-megabase rate) in specimens from each site (teal) versus all other metastatic specimens (grey). Two-sided Mann-Whitney. **(C)** Pairwise co-alteration among the 20 drivers under a null matched only on specimen number, in lung-derived specimens ($n = 83$), other metastatic specimens ($n = 842$), and their difference. Color, log odds ratio; dots, nominal $P < 0.05$. **(D)** Permutation $P$ values for lung-derived versus other metastatic specimens. dashed line, $P = 0.05$; n, lung specimens retained. **(E)** Hallmark co-activation matrices in primary tumors and lung metastases (GSE110590; Siegel et al., 2018). The difference is shown in main Fig. 1G.

Supplementary Fig. 2

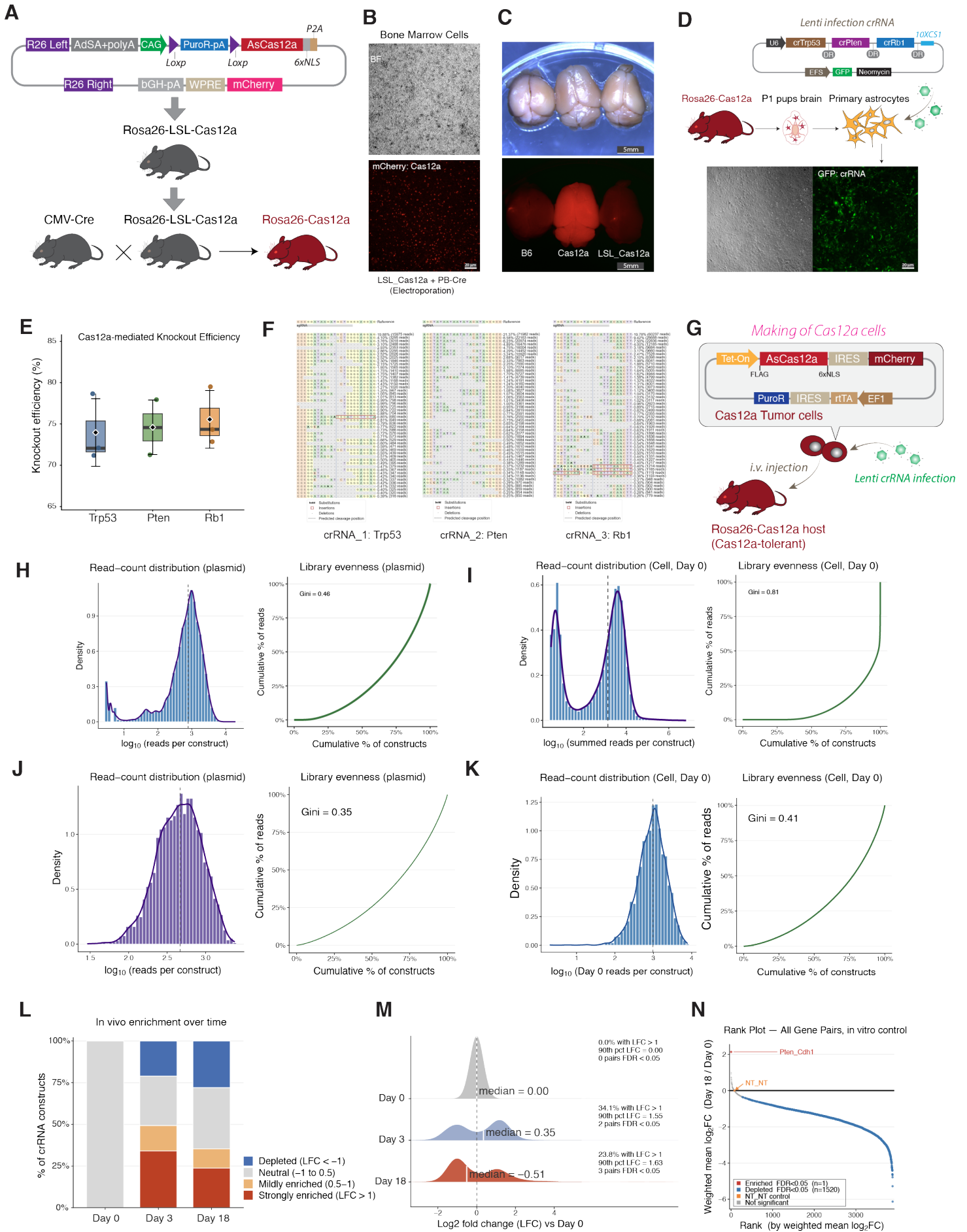

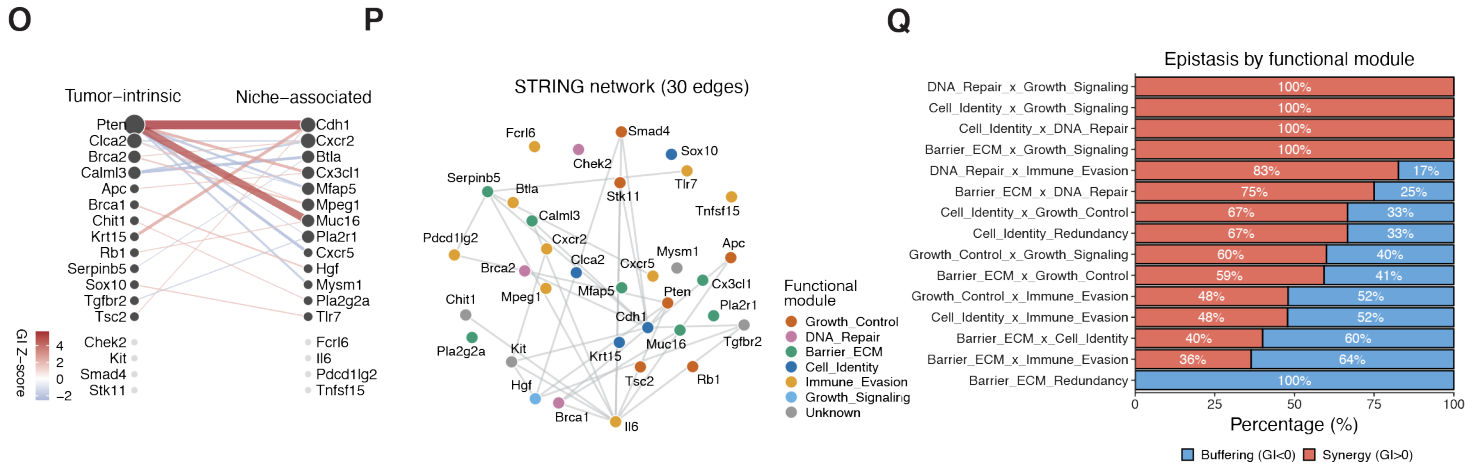

**Fig. S2. Validation of the in vivo dual-Cas12a system and quality control of the combinatorial screen (related to Fig. 2).** (A) Cre-activatable Rosa26-LSL-Cas12a knock-in allele, in which a CAG-driven PuroR-polyA cassette flanked by loxP sites blocks expression of AsCas12a-6xNLS and a P2A-linked mCherry reporter until Cre-mediated excision. Crossing to CMV-Cre yields constitutively Cas12a-expressing Rosa26-Cas12a mice. (B) mCherry reporter expression in bone marrow cells from Rosa26-LSL-Cas12a mice after PiggyBac-Cre electroporation; brightfield (top) and mCherry (bottom). Scale bar, 20 μm. (C) Whole-brain mCherry fluorescence in wild-type B6, Rosa26-Cas12a and un-recombined Rosa26-LSL-Cas12a animals, confirming that nuclease and reporter expression are Cre-dependent. Scale bars, 5 mm. (D) Lentiviral dual-crRNA vector and GFP expression in primary glia isolated from P1 Rosa26-Cas12a pups, confirming transduction of the crRNA array. Scale bar, 20 μm. (E) Cas12a-mediated knockout efficiency at Trp53, Pten and Rb1, measured by amplicon sequencing (n = 3 independent replicates per gene). Boxes span the interquartile range, whiskers the range, horizontal line marks the median and diamond the mean. (F) CRISPResso2 allele tables for the three validation crRNAs, showing substitutions, insertions and deletions relative to the reference around the predicted cleavage position. (G) Doxycycline-inducible AsCas12a system in E0771 tumor cells, carrying Tet-On AsCas12a-IRES-mCherry together with PuroR-IRES-rTA, transduced with the lentiviral crRNA library and injected intravenously into Cas12a-expressing Rosa26-Cas12a hosts. (H) Plasmid-library QC for the first-round dual-crRNA pool: read-count distribution (left) and Lorenz curve of construct evenness (right). Of the 233,684 designed constructs, 122,592 (52.5%) carry at least three reads, covering 7,097 of the 7,098 designed gene pairs and all 169 single-knockout controls, with a median of 783 reads per construct and a Gini coefficient of 0.46. (I) Injected cell-pool QC for the same first-round library, sequenced as three independent biological replicates: read-count distribution of the summed reads (left) and Lorenz curve (right). 50,990 constructs (21.8% of the design) carry at least three reads in at least one replicate, covering 7,033 gene pairs, with a median of 1,384 reads per construct and a Gini coefficient of 0.81. Replicate depths 92.7 / 92.7 / 92.5 M reads; pairwise Pearson r on $\log_{10}(\text{CPM} + 1)$, 0.79–0.87. (J) Plasmid-library QC for the second-round focused dual-crRNA pool: read-count distribution (left) and Lorenz curve of construct evenness (right). All 3,600 designed constructs are detected, with a median of 466 reads per construct and a Gini coefficient of 0.35. (K) Injected day-0 library QC for the same second-round focused pool after transduction into E0771 cells: read-count distribution (left) and Lorenz curve of construct evenness (right). Of 3,620 constructs, 3,599 (99.4%) are detected, with a median of 977 reads per construct and a Gini coefficient of 0.41. (L) Proportion of second-round crRNA constructs in each enrichment category, computed as $\log_2$ fold change relative to day 0, at day 0 (100% neutral by construction), day 3 and day 18. (M) Kernel-density ridgeline of the same distribution, one density per timepoint, annotated with the median LFC, the fraction of constructs with LFC > 1, the 90th-percentile LFC and the number of pairs surviving BH-FDR < 0.05. (N) In vitro specificity control for the first-round combinatorial CRISPR screen. The same first-round dual-crRNA library and injected-pool baseline used in (D) were cultured in vitro for 18 days. Gene pairs are ranked by weighted mean $\log_2$ fold change (Day 18/Day 0). Red and blue indicate significantly enriched and depleted gene pairs, respectively (BH-FDR < 0.05), whereas gray indicates non-significant pairs. The NT\_NT control is shown in orange. (O) Cross-program GI network in bipartite layout. The 34 second-round genes are arranged by library position. Edges connect pairs at $|GI| ZI > 1.5$, colored by GI Z-score (red positive, blue negative) with width proportional to magnitude; 23 edges among 26 connected genes. Eight genes with no strong cross-program interaction are shown as grey dots. (P) STRING protein-interaction network of the same 34 genes, plotted on the same Fruchterman-Reingold layout as main panel (I). Edges connect cross-program gene pairs at STRING medium confidence (combined score $\geq 0.40$); 30 edges. Nodes are colored by curated functional module. (Q) Epistasis direction by curated functional module combination. For each pair of modules (excluding Unknown), the bar shows the proportion of gene pairs classified as synergistic (GI > 0, red) or buffering (GI < 0, blue); 15 module combinations spanning 189 annotated pairs.

Supplementary Fig. 3

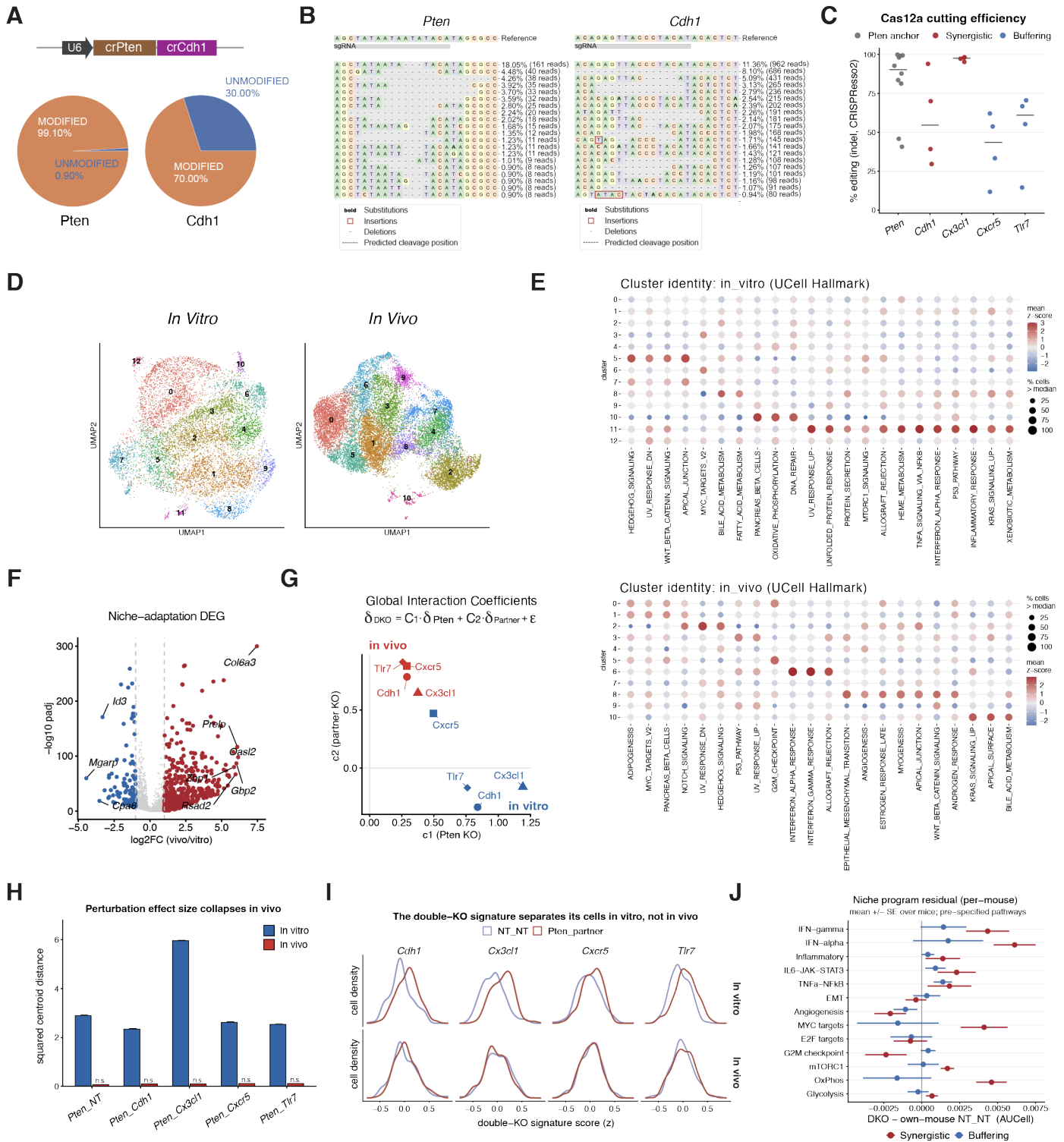

**Fig. S3. Metastatic niche selection determines Pten-partner epistasis outcome (related to Fig. 3).** (A) Structure of a representative dual-crRNA construct (U6-crPten-crCdh1) and CRISPResso2 quantification of editing at the two target sites in the edited line, showing 99.10% modified reads at Pten and 70.00% at Cdh1. (B) CRISPResso2 allele plots at the Pten and Cdh1 target sites, giving the most frequent alleles with their percentage and read count. Substitutions are shown in bold, insertions boxed and deletions dashed; the dotted line marks the predicted cleavage position. (C) AsCas12a editing efficiency by CRISPResso2 for each target gene across the dual-guide constructs. Each point is one construct or sample and the crossbar is the median: Pten 90.2% (n = 10), Cdh1 54.6%, Cx3cl1 97.6%, Cxcr5 43.5% and Tlr7 61.0% (n = 4 each). Color denotes epistasis class. (D) UMAP of all tumor cells with cluster identity, in vitro (13 clusters) and in vivo (11 clusters). (E) Cluster identity as UCell Hallmark enrichment for the in vitro and in vivo clusters. Fill is the mean score z-scored across clusters and size is the percentage of the cluster's cells scoring above the context-wide median for that program. (F) Differential expression between the in vivo and in vitro NT\_NT control: 830 genes are higher in vivo and 131 higher in vitro at a Benjamini-Hochberg threshold of 0.05, including Col6a3, Preip, Gbp2, Oasl2 and Rsad2 among those induced and Id3 and Mgar among those reduced. (G) Global interaction coefficients from the additive epistasis model $\delta_{DKO} = c_1 \cdot \delta_{Pten} + c_2 \cdot \delta_{partner} + \epsilon$. Fitted coefficients for each Pten-partner pair are shown for in vivo (red) and in vitro (blue) conditions, with $c_1$ (Pten KO coefficient) and $c_2$ (partner KO coefficient) plotted on the x- and y-axes, respectively. Symbol shapes denote individual partner gene. (H) Squared centroid distance of each perturbation from NT\_NT in z-scored principal-component space, in vitro versus in vivo. Effect sizes collapse in vivo and none remains significant. This statistic and its null differ from those of Fig. 3F and the two panels are not numerically comparable. (I) Density of the double-knockout signature score in NT\_NT and in Pten\_partner cells for each partner, in vitro above and in vivo below. The signature separates the two genotypes in vitro but not in vivo. (J) Per-mouse niche-program residual across the 13 pre-specified pathways, as mean ± s.e.m. over mice (synergistic n = 3, buffering n = 4). No pathway separates the two classes at the animal level (all Benjamini-Hochberg $q \geq 0.78$).

Supplementary Fig. 4

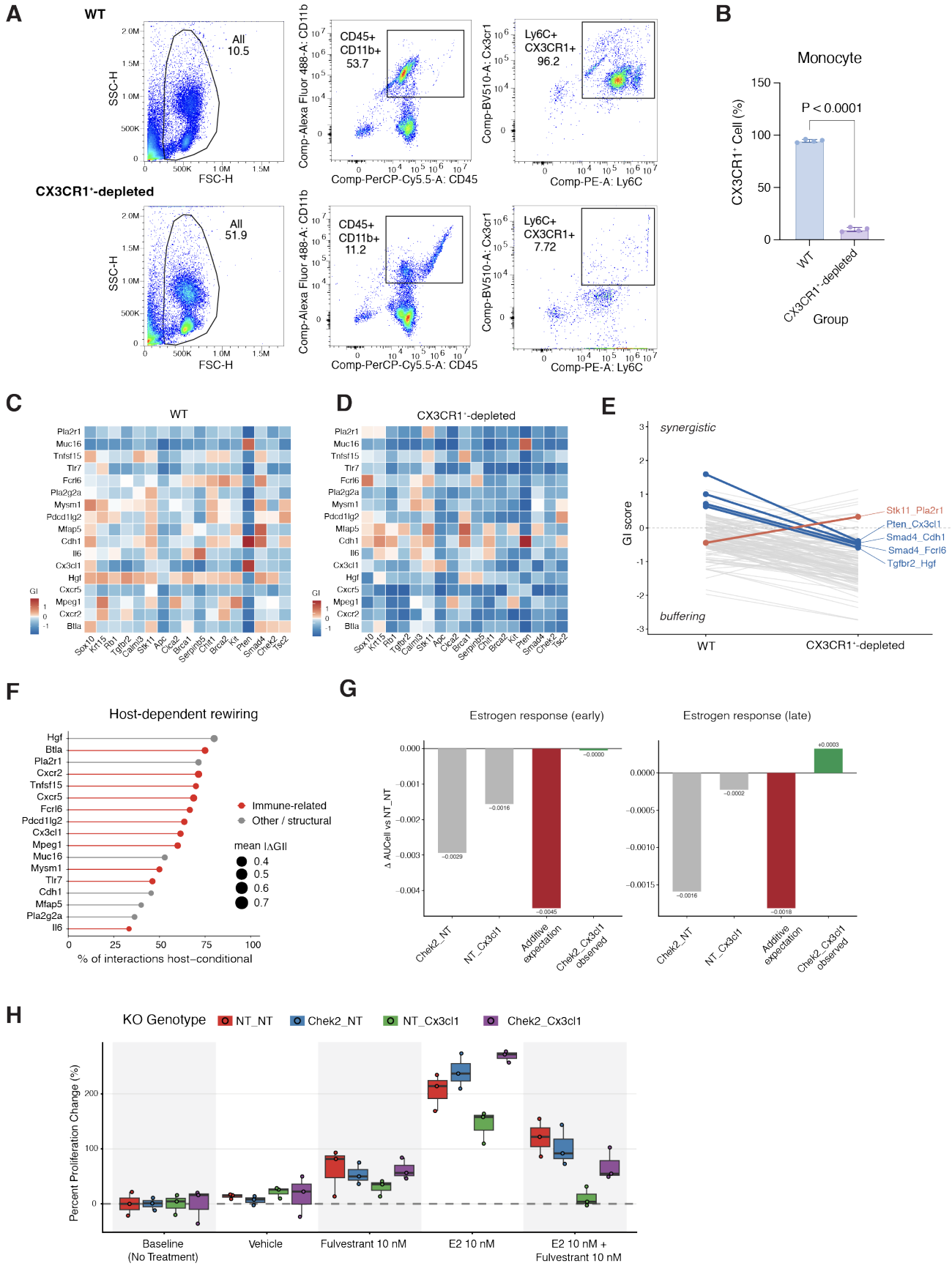

**Fig. S4. CX3CR1<sup>+</sup> myeloid-cell depletion, host-specific genetic interaction maps, and validation of host-dependent interactions (related to Fig. 4).** (A) Representative flow-cytometry gating of the CX3CR1<sup>+</sup> myeloid compartment in WT (top) and CX3CR1<sup>-</sup>-depleted (bottom) hosts. Cells were sequentially gated by scatter, CD45<sup>+</sup>CD11b<sup>+</sup> expression, and Ly6C<sup>+</sup>CX3CR1<sup>+</sup> expression. Percentages indicate the fraction of cells within the corresponding parent gate. (B) Quantification of CX3CR1<sup>+</sup> cells among CD45<sup>+</sup>CD11b<sup>+</sup> cells in WT and CX3CR1<sup>-</sup>-depleted hosts. Bars indicate mean values and points represent individual mice. Statistical significance was assessed using Welch's two-sample t-test; $P < 0.0001$. (C, D) GI scores in WT (C) and CX3CR1<sup>-</sup>-depleted (D) hosts. Gene order and matrix orientation are identical between panels and correspond to those in Fig. 4B, with rows and columns ordered by hierarchical clustering of the ΔGI matrix. The diverging color scale is centered at zero. \*$q < 0.05$; $q < 0.10$. (E) GI scores for interactions exhibiting significant sign reversal between WT and CX3CR1<sup>-</sup>-depleted hosts, defined as interactions with $q < 0.05$ in both host conditions and opposite GI signs. Colored lines indicate sign-reversing interactions; gray lines indicate other host-conditional interactions. (F) Host-conditional interaction rate for each niche-associated partner gene, calculated as the fraction of tested driver-partner interactions with significant ΔGI ($q < 0.05$). Point size represents the mean $|\Delta$GI across interactions, and color denotes partner category (immune-related versus other/structural). (G) Estrogen-response pathway scores for Chek2\_Cx3cl1 genotypes measured by Perturb-seq. For the early and late estrogen-response programs, bars show the change in AUC score relative to NT\_NT for Chek2 and Cx3cl1 single knockouts, the additive expectation derived from the two single knockouts, and the observed Chek2\_Cx3cl1 double knockout. (H) Proliferation responses of Chek2\_Cx3cl1 genotypes to estrogen-receptor modulation. E0771 cells carrying NT\_NT, Chek2\_NT, NT\_Cx3cl1, or Chek2\_Cx3cl1 perturbations were assayed under baseline (no treatment), vehicle, fulvestrant (10 nM), 17β-estradiol (E2; 10 nM), or combined E2 and fulvestrant (10 nM each) conditions. Percent change in proliferation is shown; boxes indicate the interquartile range and points represent biological replicates ($n = 3$).

Supplementary Fig. 5

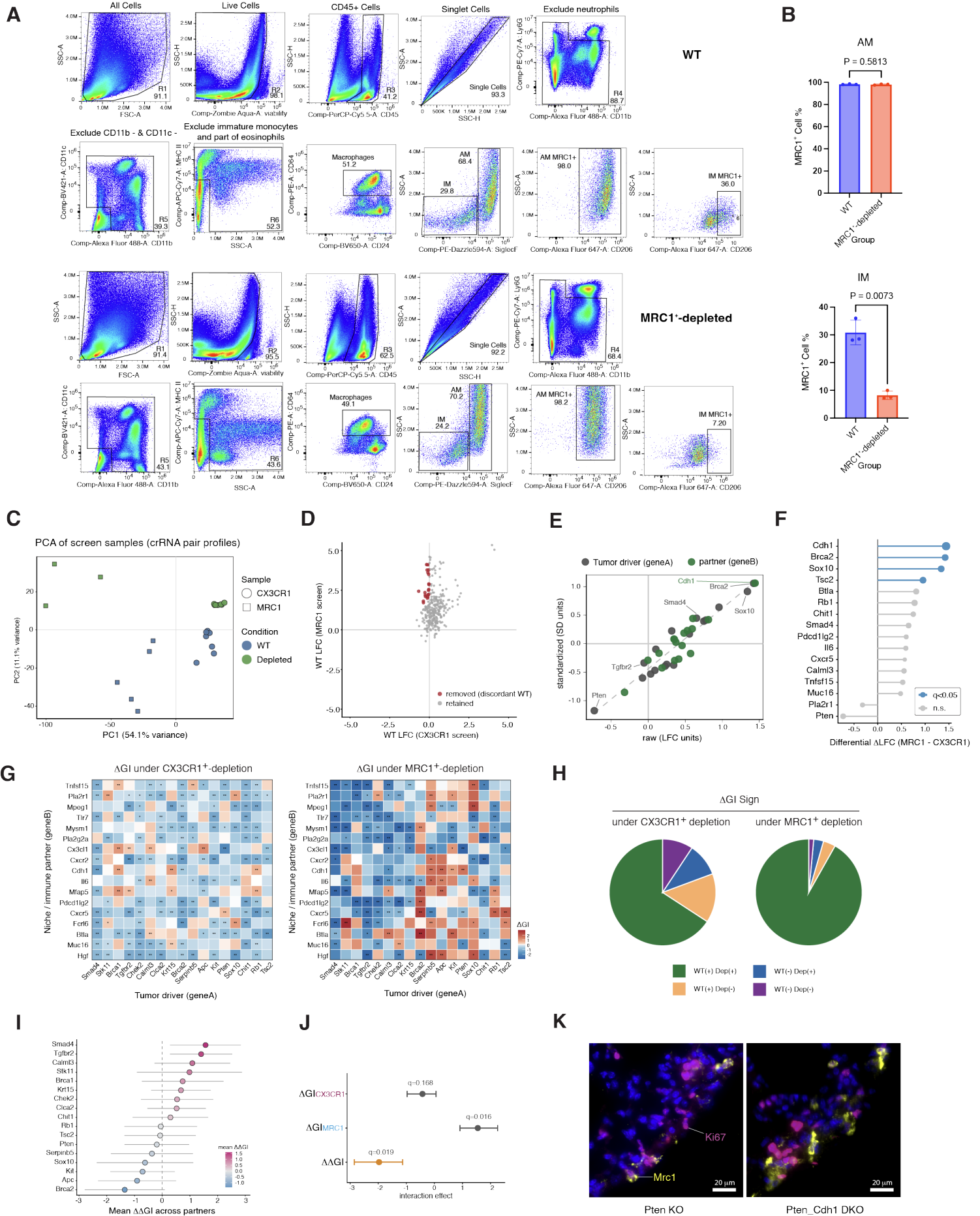

**Fig. S5. MRC1<sup>+</sup> macrophage depletion and comparative analysis of niche-dependent genetic interaction rewiring (related to Fig. 5).** (A) Representative flow-cytometry gating of lung macrophage populations in WT (top) and MRC1<sup>-/-</sup>-depleted (bottom) hosts. Live CD45<sup>+</sup> singlets were sequentially gated to exclude Ly6G<sup>+</sup> neutrophils, CD11b<sup>+</sup>CD11c<sup>+</sup> cells, and MHC II-defined immature monocytes and eosinophils. Macrophages were identified as CD64<sup>+</sup>CD24<sup>+</sup> cells and separated into alveolar (AM) and interstitial (IM) macrophage populations by SiglecF expression. MRC1<sup>+</sup> cells within each population were identified by CD206 staining. Percentages indicate the fraction of cells within the corresponding parent gate. (B) Quantification of MRC1<sup>+</sup> cells among alveolar (AM) and interstitial (IM) macrophages in WT and MRC1<sup>-/-</sup>-depleted hosts. Bars indicate mean ± SD and points represent individual mice (n = 3 per group). Statistical significance was assessed using an unpaired two-sample t-test; P values are indicated. (C) Principal component analysis (PCA) of sample-level crRNA-pair abundance profiles from the CX3CR1<sup>-/-</sup> and MRC1<sup>-/-</sup>-depletion screens. Samples are distinguished by screening model (circles, CX3CR1<sup>-/-</sup>; squares, MRC1<sup>-/-</sup>) and host condition (blue, WT; green, depleted). Percent variance explained by each principal component is indicated on the axes. (D) Cross-screen comparison used to identify discordant WT measurements. LFC values for all 289 gene pairs in WT hosts from the MRC1 screen are plotted against those from the CX3CR1 screen. Pairs with opposite-signed WT effects and an absolute between-screen difference >2 were excluded (red); retained pairs are shown in gray. (E) Comparison of gene-level first-order effects calculated from raw LFC values and screen-standardized LFC values. Tumor-driver genes (gene A) and partner genes (gene B) are indicated in gray and green, respectively; the dashed line denotes the identity line. (F) Differential first-order effects between the MRC1<sup>-/-</sup> and CX3CR1<sup>-/-</sup>-depletion screens ($\Delta$ LFC, MRC1 – $\Delta$ LFC, CX3CR1) for individual genes. Genes are ordered by differential $\Delta$ LFC; blue indicates $q < 0.05$ and gray indicates non-significant differences. (G) Host-dependent changes in genetic interactions following CX3CR1<sup>-/-</sup> (left) or MRC1<sup>-/-</sup> (right) cell depletion. ΔGI was calculated as GI<sub>depleted</sub> – GI<sub>WT</sub> within each screen. Tumor-driver genes are shown in columns and niche/immune partner genes in rows. Asterisks indicate statistically significant ΔGI values based on bootstrap analysis. (H) Distribution of GI sign combinations between WT and depleted hosts for the CX3CR1<sup>-/-</sup> and MRC1<sup>-/-</sup>-depletion screens among the 269 gene pairs retained after WT consistency filtering. Interactions are classified according to their GI signs in WT and depleted conditions. (I) Driver-level summary of differential niche dependence. For each tumor-driver gene, $\Delta\Delta$GI values between the MRC1<sup>-/-</sup> and CX3CR1<sup>-/-</sup>-depletion screens were summarized across its partner genes. Points indicate the mean $\Delta\Delta$GI and error bars indicate SD. (J) Host-dependent interaction effects for Pten\_Cdh1. ΔGI values following CX3CR1<sup>-/-</sup> and MRC1<sup>-/-</sup>-depletion and the corresponding between-screen $\Delta\Delta$GI are shown with 95% bootstrap confidence intervals. The dashed vertical line indicates zero interaction effect; q values are indicated. (K) Representative immunofluorescence images of lung metastatic lesions formed by Pten single-knockout (Pten KO) or Pten\_Cdh1 double-knockout (Pten\_Cdh1 DKO) E0771 cells. Tumor lesions were identified by tumor-cell GFP; Ki67 is shown in magenta, MRC1/CD206 in yellow, and nuclei in blue. Scale bars, 20 μm.
